# Enkephalin constrains fear learning via volume transmission to the lateral amygdala

**DOI:** 10.64898/2026.07.17.739254

**Authors:** Roger Wang, Joanna Yau, Muskaan Kalra, Sophia Gilchrist, Abbey Livermore, Gabrielle Gregoriou, Merle Schennink, Cesar Moreno, Rakulan Santhakumar, Neda Rafiei, Zian Hao, Julianne Estrella, Thea Thomsen, Emilie Hartvig, Maria Capertonova, Katja Jensen, Sidsel Landler, Lin Tian, Greg Neely, Andre Berndt, Michael Bruchas, Gavan McNally, Elena Bagley

## Abstract

Fear learning involves the formation of associations between cues and aversive outcomes, a process that must be tightly regulated to prevent excessive or generalised fear. Dopamine release in the lateral amygdala (LA) drives fear acquisition, whereas endogenous opioids constrain it. However, whether opioids are dynamically released within the amygdala circuits during learning, and how they exert this control remain unclear. Here we show that met-enkephalin is locally released within the amygdala during auditory fear conditioning, with signals shifting from the aversive outcome to its predictive cue as learning progresses. The amygdalo-striatal transition zone (ASt), is the principal source of this enkephalin, released from medium spiny neurons receiving strong auditory thalamic input. This enkephalin spreads from the ASt to the LA via volume transmission. Selective knockdown in the ASt abolished opioid signals and enhanced fear learning, demonstrating that this diffuse signal constrains fear memory formation. We further show that enkephalin suppresses dopamine release in both the ASt and LA via µ-opioid receptors, identifying the ASt as a neuromodulator hub coordinating opioid and dopaminergic signalling across amygdala fear circuits. Although demonstrated here for auditory fear learning, the ASt receives multimodal sensory input, suggesting a broader mechanism through which sensory experience recruits enkephalin release to gate associative learning

## Introduction

Fear learning allows organisms to predict and respond to environmental threats by forming associations between neutral cues and aversive outcomes. To be adaptive, these associations must be appropriately scaled, insufficient learning risks failure to avoid danger, whereas excessive or generalized fear contributes to anxiety-and trauma-related disorders [1–3]. Understanding the mechanisms that both drive and constrain fear learning is therefore essential for identifying how fear memories form and how they may be controlled.

Fear learning depends on synaptic plasticity within the amygdala circuits that link sensory and aversive information [4–9]. Dopamine plays a central role in this process [4, 10, 11]. Dopaminergic projections from the ventral tegmental area (VTA) to the lateral amygdala (LA) promotes fear acquisition by facilitating synaptic plasticity at sensory inputs [4, 11], whereas reducing dopamine signalling in these circuits impairs fear learning [11–13]. These findings support a role for dopamine as a teaching signal that strengthens cue-threat associations.

If dopamine promotes fear learning, then a complementary inhibitory mechanism is required to prevent excessive fear acquisition. Activation of µ-opioid receptors attenuates fear learning in rodents [14], whereas opioid receptors blockade enhances fear acquisition and second-order conditioning [15, 16], indicating that endogenous opioids actively constrain associative learning [17]. In humans, the μ-opioid receptor agonist buprenorphine reduces fearful responding [18], and antagonist enhances acquisition of conditioned fear [19, 20], establishing that the µ-opioid receptor mediated constraint is conserved across species.

Despite this well-established role, the specific circuits through which endogenous opioids limit fear learning remain poorly understood. The amygdalo-striatal transition zone (ASt), a region adjacent to the LA is enriched with the endogenous opioid enkephalin [21, 22] and receives strong dopaminergic input from the VTA [11, 23]. VTA neurons projecting to the amygdala and striatum also represent distinct subpopulation with different opioid receptor profiles [24]. The ASt, a region containing neuronal population similar to striatum [25], and sits at the interface of these two regions, is therefore particularly important site for understanding opioid-dopamine interactions in fear learning.

Here we show that enkephalin is released within the amygdala during fear conditioning, originating from enkephalinergic medium spiny neurons of the ASt which can be driven by auditory thalamic activity, and spreads to the LA via volume transmission where it suppresses dopamine release via µ-opioid receptor activation. These findings identify the ASt as a neuromodulator hub in which sensory-driven enkephalin release shapes dopaminergic signalling across amygdala circuits to constrain fear learning.

## Results

### Opioids signals track fear learning in the amygdala

Prior *in vivo* pharmacological studies suggests that endogenous opioid acts through µ-opioid receptors to limit fear learning [15, 17, 20]. Amongst the endogenous opioid peptides with high affinity for µ-opioid receptors [26], enkephalin is expressed locally in the amygdala and surrounding regions [22] whereas β-endorphin is primarily expressed in arcuate nucleus of the hypothalamus neurons which provide sparse innervation of the lateral amygdala [27], making enkephalin the more likely candidate for mediating local opioid signalling. We therefore chose δLight1.1, a genetically encoded sensor based on the δ-opioid receptors, because enkephalin binds both µ-and δ- receptors with high affinity [26] and δlight1.1 has the highest sensitivity to enkephalin amongst the endogenous opioid peptides [28], making it the best tool to monitor enkephalin release dynamics *in vivo*.

To determine whether endogenous opioids are dynamically released and involved in fear learning, we expressed δLight1.1 (*n =* 11) in the lateral amygdala (LA) and monitored fluorescent changes during auditory fear conditioning (Fig 1. A, B). Rats underwent lever press training before conditioned fear training, in which a 60 s auditory tone (CS) co-terminated with a 0.5 s footshock (US). Conditioned suppression of lever pressing was used as the measure of fear, with suppression ratios approaching zero indicating strong fear. Animals showed robust fear acquisition, with suppression ratios progressively decreasing across the training (Fig 1. D), indicating that the CS acquired predictive value for the aversive outcome.

**Figure 1.**
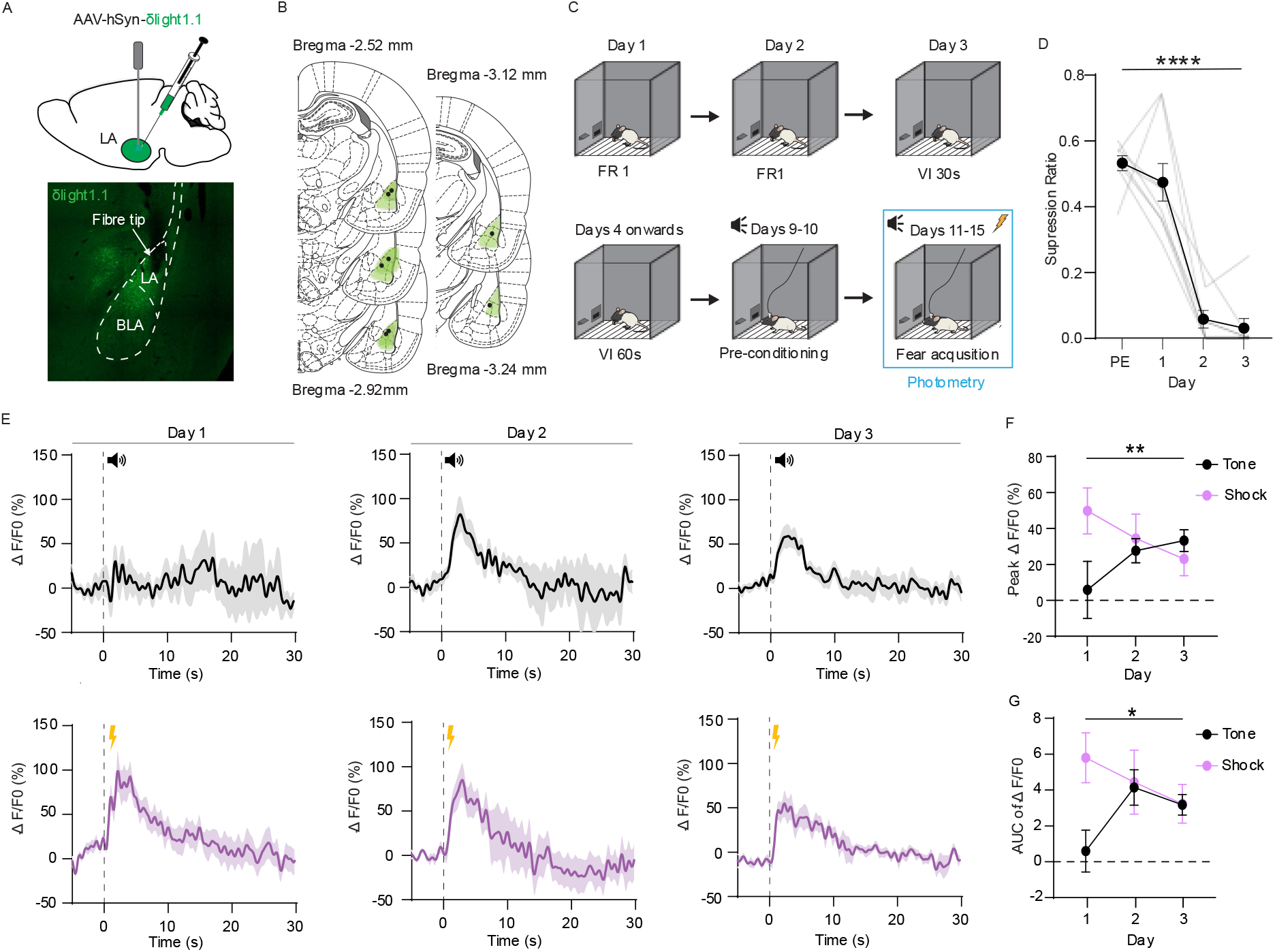
Endogenous opioids released in the amygdala exhibit prediction error-like dynamics during fear conditioning. **(A)** Top: Schematic diagram indicating AAV9-hSyn-δlight1.1 injection into the lateral amygdala (LA) and an optical fibre implanted above the injection site. Bottom: Representative images showing fluorescence of δLight1.1 with fibre placement. **(B)** Schematic representation of LA δLight1.1 expression and optical fibre tips for all animals. **(C)** Cartoon of the behavioural protocol. Rats first learnt to lever press for reward under fixed ratio 1 (FR1) then maintained on variable interval 30s (VI 30s) and variable interval 60s (VI 60s). Rats are then exposed to 60s auditory tone (CS) co-terminating with 0.5s shock (US), during which fibre photometry recordings are made. **(D)** Suppression ratios during pre-exposure (PE) and across training days. days (one-way repeated measure ANOVA, linear trend across days, *F* _(1,21)_ = 151.1, P < 0.0001). **(E)** Average δLight1.1 fluorescent signals for tone (black) and shock (purple) across training days. *n* = 11. **(F)** Chart showing peak ΔF/F_0_ responses measured as an average of the first 5 seconds post tone (black) and shock (purple). linear regression, Tone: R^2^ = 0.11, β = 0.14, 95% confidence interval-1.31 to 28.76; Shock R^2^ = 0.7, β =-13.37, 95% confidence interval-30.48 to 3.75). Data represented as mean ± SEM. **(G)** Area under the δLight1.1 curve for tone and shock across training days (linear regression, Tone: R^2^ = 0.1, β = 0.51, 95% confidence interval-0.05 to 1.08; Shock R^2^ = 0.9, β =-2.04, 95% confidence interval-2.3 to-1.8) Data presented as mean ± s.e.m with individual data points. *P<0.05,****P<0.0001

Photometry recordings revealed that early in training, CS presentations produced minimal δLight fluorescent changes, whereas the US evoked a robust, time-locked increase in fluorescent signals (Fig 1. E), indicating increased extracellular concentrations of endogenous opioids. However, as the CS became increasingly predictive of the US across sessions, the δLight responses transferred from the outcome (US) to the predictive cue (CS) (Fig 1. E). Both CS- and US- evoked opioid signals had rapids onsets and lasted approximately 10 seconds, consistent with a transient peptide release event at stimulus onset. Quantification of response magnitude via area under the δLight1.1 curve (AUC) demonstrated that US-evoked signals decreased across training, reflected in a negative slope (β =-2.04, 95% confidence interval-2.3 to-1.8), whereas CS-evoked signals increased, exhibiting a positive slope (β = 0.53, 95% confidence interval-0.05 to 1.08). These slopes differed significantly from each other (F _(1,62)_ = 72.32, P < 0.0001, Fig. 1 F). Thus, endogenous opioids are dynamically released in the LA during fear learning and its signalling shifts from the aversive outcomes to its predictive cue across training.

### Met-enkephalin is the dominant endogenous opioid released from the ASt

The LA itself is relatively sparse in endogenous opioid expression [27], raising the question of where the opioid signals observed during fear learning originate. The ASt, situated immediately adjacent to the LA, expresses high levels of the endogenous opioid enkephalin and dynorphin within medium spiny neurons [29] and receives convergent sensory and dopaminergic input [23, 30], positioning it as a likely local source of endogenous opioid release during fear learning.

We first expressed δLight1.1 in the ASt to monitor fluorescence changes in acute brain slices (Fig. 2 A). Bath application of Met-enkephalin produced robust increases in fluorescence that were reversed by naloxone (Fig 2. B, C), demonstrating reliable detection of extracellular opioid concentration. To identify which endogenous opioid activates δLight1.1, we established concentration-response curves for met-enkephalin, dynorphin and β-endorphin. Met-enkephalin produced a robust concentration-dependent increase in fluorescence (Fig 2. D, E). dynorphin and beta-endorphin required substantially higher concentrations to produce comparable responses (Fig 2. E), consistent with the preferential sensitivity δLight1.1 for enkephalin [28]. Therefore, the relative sensitivity of δLight1.1 can be used to infer identity of the released peptide from the amplitude of evoked signals.

**Figure 2.**
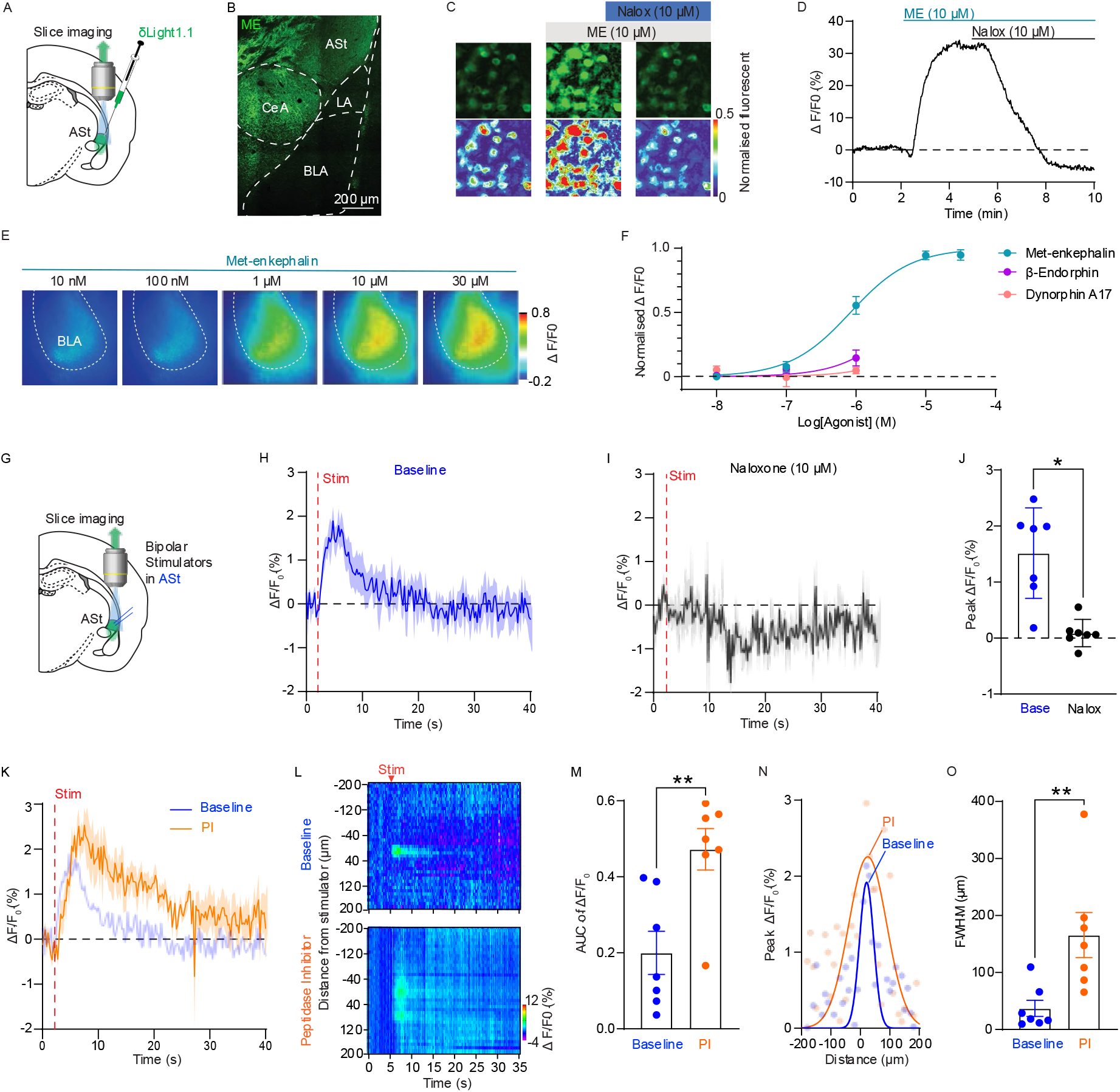
Met-enkephalin is the predominant endogenous opioid released from the amygdalo-striatal transition zone. **(A)** Schematic diagram showing δLight1.1 injection sites. **(B)** Pseudo-coloured δLight1.1 images and heatmap showing increases in fluorescence following bath application of met-enkephalin (10 µM)) and reversal by naloxone (10 µM) **(C)** Representative trace showing ΔF/F_0_ changes over time during met-enkephalin and naloxone application. **(D)** Heatmaps showing ΔF/F_0_ changes for increasing concentrations of met-enkephalin. **(E)** Normalised ΔF/F_0_ concentration-response curves for met-enkephalin, dynorphin A17 and β-endorphin (non-linear regression, unconstrained fit). **(F)** Schematic diagram showing imaging and bipolar electrode placement in the ASt. **(G-H)** Average traces showing ΔF/F_0_ changes following electrical stimulation of the ASt (100 pulses at 50 Hz) during baseline and naloxone application. **(I)** Chart showing peak ΔF/F_0_ responses measured as an average of the first 8 frames post-stimulation in the ASt during baseline and naloxone application. Data presented as mean ± sem with individual data points. (P < 0.01, Paired Student’s t-test, baseline versus naloxone). **(J)** Average traces showing ΔF/F_0_ changes following electrical stimulation of the ASt during peptidase inhibitors application. Shade blue trace represents baseline from (G). **(K)** Heatmaps showing the spatial-temporal profile of evoked fluorescence during baseline and peptidase inhibitor application, averaged across slices, individual dots represent ΔF/F_0_ as measured at specific distances. **(L)** Area under the δLight1.1 curve for baseline and peptidase inhibitor application in the ASt. (P < 0.01, Paired Student’s t-test, baseline versus peptidase inhibitor). **(M)** Average spatial profile of evoked fluorescence measured at 2 s post-stimulation during baseline and peptidase inhibitor application, each fitted with a Gaussian function. Opaque dots represent averaged ΔF/F_0_ measured each distance. **(N)** Full width at half maximum (FWHM) for baseline versus peptidase inhibitor application in the ASt. (P < 0.01, Paired Student’s t-test, baseline versus peptidase inhibitor).

We next asked whether electrical stimulation of the ASt is sufficient to evoke endogenous opioid release locally (Fig 2. F). Local electrical stimulation of the ASt evoked robust fluorescence signals (Fig 2. F-I), that were abolished by naloxone (Fig 2. F-I), indicating that these signals reflected endogenous opioid binding to the sensors. Given that δLight1.1 requires substantially higher concentration of dynorphin and β-endorphin to produce comparable signals, the amplitude of these responses is most consistent with the stimulation triggering met-enkephalin release. Peptidase inhibitor application (10 μM Bestatin, 10 μM Thiorphan and 1 μM Captopril) did not increase the peak responses evoked by electrical stimulation (Baseline: 1.515±0.26% ΔF/F_0_, PI: 1.296±0.36 ΔF/F_0_, *n = 7*, P = 0.72, Paired Student’s t-test, baseline versus PI, Fig 2 J) but significantly prolonged response duration with a > 2-fold increase in the AUC (Fig 2. J, L). Peptidase inhibition also substantially expanded the spatial extent of the signal. Quantification using full width at half maximum (FWHM) measurements demonstrated a significant increase in enkephalin spread in the ASt (Fig 2. M, N), consistent with enzymatic degradation normally acting to limit the spatial reach of the enkephalin rather than its peak amplitude. The selective enhancement by peptidase inhibitors which preferentially target enkephalin over other opioid peptides [31, 32] further supports met-enkephalin as the principal released peptide, and the spatial expansion indicates that peptidase actively constrains the spread of enkephalin through the extracellular space.

In the ASt enkephalin is only expressed in D2 containing MSN’s [29], suggesting these neurons are the primary source of enkephalin in the LA. If so, then D2 receptor activation should reduce enkephalin release. To test this, we expressed δLight1.1 in the ASt and bath applied selective D2 agonist quinpirole and antagonist raclopride and monitored electrically evoked δLight signals. Bath application of the selective D2 agonist quinpirole (10 μM) suppressed δLight signals evoked by local electrical stimulation (Baseline: 2.304±0.89% ΔF/F_0_, quinpirole: 0.17±1.20% ΔF/F_0_, *n = 6*, P = 0.009, Paired Student’s t-test, baseline versus quinpirole, Supp Fig 1). This inhibition was reversed by the D2 selective antagonist raclopride (Raclopride: 1.90±0.25% ΔF/F_0_, *n = 6*, P = 0.02, Paired Student’s t-test, quinpirole versus raclopride, Supp Fig 2). Together these data indicate that D2-expressing medium spiny neurons are a principal source of enkephalin release in the ASt.

These converging lines of evidence indicate that met-enkephalin released from D2-expressing MSNs is the dominant endogenous opioid released in amygdala fear circuits and identify the ASt as the likely source of the opioid signals observed in the LA during fear learning.

### Auditory thalamic inputs trigger enkephalin release from the ASt that diffuses to LA

We next asked what drives enkephalin release from the ASt during auditory fear learning and whether this signal diffuses into the LA. The medial geniculate nucleus (MGN) of the auditory thalamus is the primary relay of auditory information to the amygdala and MGN projections to the LA are critical for fear memory acquisition [6, 8, 33]. Interestingly, the MGN also sends dense projections to the ASt [33], prompting us to examine whether auditory thalamic activity could drive enkephalin release. We expressed AAV-ChR2 into the MGN and performed whole-cell recordings in both the LA and ASt (Fig 3. A). ChR2 expression was observed in MGN cell bodied with dense terminal labelling in the ASt (Fig 3. B) and comparatively sparser terminal labelling in the LA (Fig 3 B). Optical stimulation of MGN terminals evoked excitatory postsynaptic currents (oEPSC’s) in both pyramidal neurons in the LA and putative medium spiny neurons in the ASt that were abolished by the AMPA/kainate receptor antagonist CNQX (Fig 3. C, D). We observed functional MGN connectivity in approximately half of the total recorded LA neurons (36.09 ± 7.08 pA, *n = 51* Fig 3. C), consistent with previous reports [34]. In contrast, recordings in the ASt revealed that all neurons received functional MGN glutamatergic input (185.3 ± 18.75 pA, *n = 44)*, with significantly larger synaptic response (two tailed unpaired *t-*test, P < 0.0001. Fig. 3. C, D), demonstrating that auditory thalamic activity preferentially engages the ASt.

**Figure 3.**
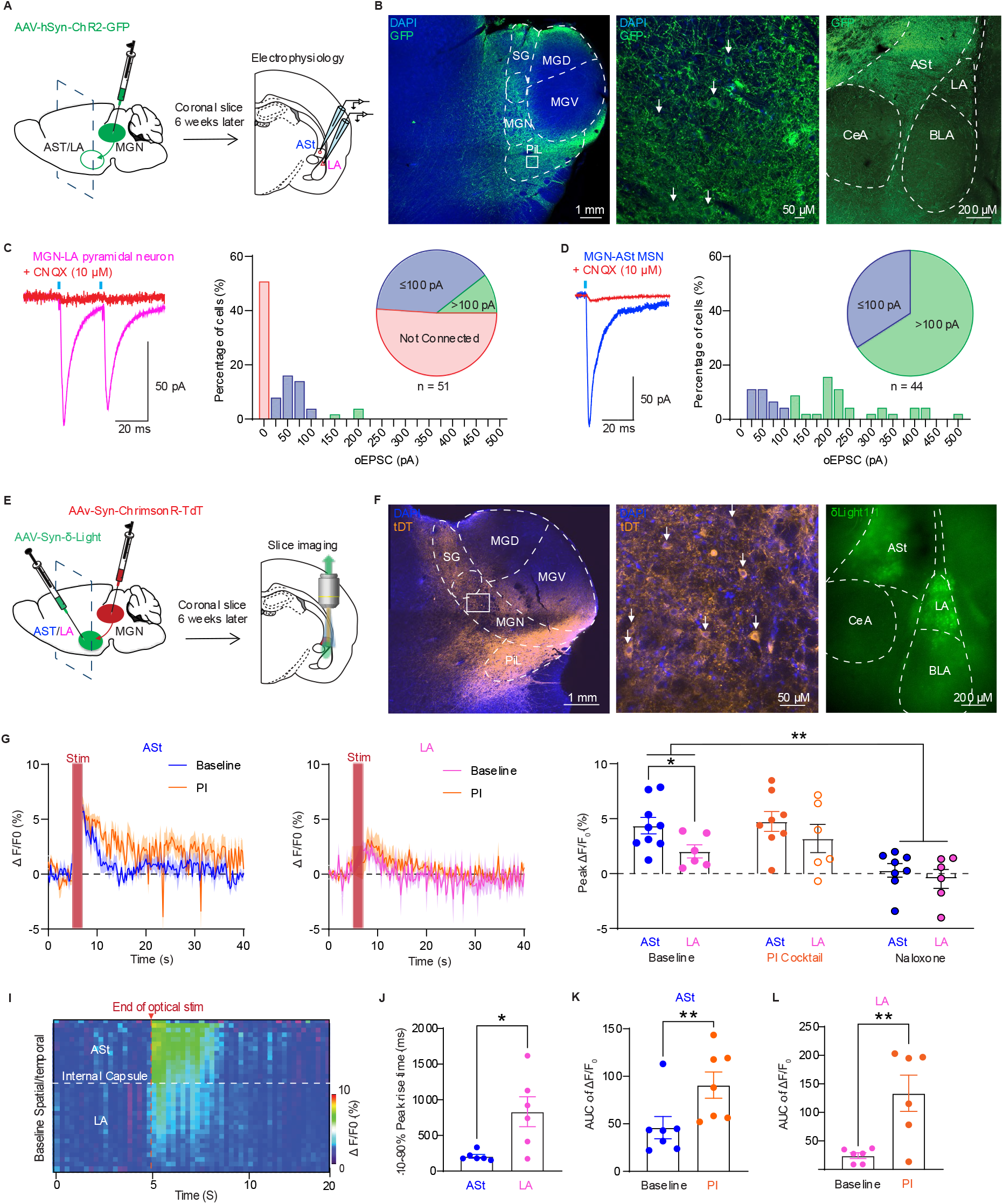
Auditory thalamic drive triggers enkephalin release from the ASt that diffuses to the lateral amygdala. **(A**) Schematic diagram showing injection of AAV-ChR2 into the MGN and recording locations in the ASt and LA. **(B)** *Post hoc* confocal images showing DAPI (blue) and GFP (green) expression in the auditory thalamus and amygdala. Magnification of white box shows expression of GFP in cell soma within the MGN. **(C)** Example oEPSCs of LA neurons (1ms optical stimulation) during baseline and CNQX application and frequency distribution of oEPSC size. **(D)** Same as in (C) but for neurons patched in the ASt (Two tailed unpaired *t-*test ASt vs LA amplitude, P < 0.0001). **(E)** Schematic diagram showing injection of AAV-ChrimsonR into the MGN and the fluorescent sensor δLight1.1 into the ASt/LA. Slices were taken 6 weeks post-injection for fluorescent recordings. **(F)** *Post hoc* confocal images showing DAPI (blue) and tdTomato (orange) expression in the auditory thalamus and pseudo-coloured live δLight1.1 (green) expression in the ASt and LA Magnification of white box shows expression of tdTomato in cell soma within the MGN. **(G)** Average traces showing ΔF/F_0_ changes evoked by optical stimulation (100 stimulations at 50 Hz) as measured in the ASt and LA during baseline and peptidase inhibitor. **(H)** Chart showing peak ΔF/F_0_ responses measured as an average of the first 8 frames post-stimulation in the ASt and LA during baseline, peptidase inhibitor and naloxone application. (Two-way repeated measures ANOVA, Sidak’s multiple comparison, Baseline ASt vs Baseline LA: P = 0.026; Baseline vs Naloxone: P = 0.005). **(I)** Heatmap showing spatial-temporal profile of the evoked fluorescence in the ASt and LA. **(J)** 10-90% rise times measured in the ASt and LA. (Two-tailed unpaired *t*-test, P = 0.032). **(K-L)** Area under the δLight1.1 curve measured from the ASt and LA. (Two-tailed paired *t-*test, ASt: P = 0.007; LA: P = 0.02). Data presented as mean ± s.e.m with individual data points

To test whether this strong thalamic input drives enkephalin release and consequently also increases extracellular enkephalin in the LA, we combined injection of the red-shifted channel rhodopsin ChrimsonR in the MGN with expression of the opioid sensor δ-light in the ASt and LA (Fig 3. E). ChrimsonR expression was confirmed in MGN cell bodies with terminal expression visible in both the ASt and LA, δ-light expression was observed throughout both regions as well (Fig 3. F). Optical stimulation of MGN terminals produced robust time-locked increases in fluorescence in the ASt that was abolished by naloxone (Fig 3 C, E). Interestingly, MGN terminal stimulation drove significantly larger responses than local electrical stimulation (Electrical: 1.515±0.81% ΔF/F_0,_ *n = 6*, Optical: 4.367±0.75% ΔF/F_0,_ *n = 9*, P = 0.006, Unpaired Student’s t-test, electrical versus optical). Notably, δ-light signals were also observed in the LA in response to optical stimulation of MGN terminals, though the peak response in the LA were significantly smaller than in the ASt (Fig 3 D, E). We reasoned that the smaller peak fluorescence seen in the LA could be due to diffusion of enkephalin across the ASt to the LA. Spatial-temporal analysis revealed that peak LA signals exhibited a significantly slower 10-90% peak rise time compared to ASt responses (ASt: 207.3 ± 26.63 ms; LA: 832.7 ± 210.6 ms; two tailed paired *t-*test, P = 0.032, Fig 3. I, J). Peptidase inhibitors significantly increased the area under the δLight curve in both the ASt (two tailed paired *t*-test, P = 0.007, Fig. 3 K) and the LA (two tailed paired *t-*test, P = 0.02, Fig 3 L), demonstrating that extracellular degradation normally constrains the persistence of enkephalin signalling, possibly more at distal sites. This suggests that alterations in peptidase activity, such as those induced by withdrawal may alter the spatial extend to which enkephalin can act [35]. Together with the smaller amplitude and delayed onset of LA signals relative to those in the ASt indicate that enkephalin released from ASt neuron spreads into the LA rather than being released locally within it.

To examine whether the anatomical organisation of the ASt could support enkephalin delivery to the LA, we filled individual ASt MSNs with biocytin and performed post-hoc immunohistochemistry for met-enkephalin (Fig S3. A). Following post-hoc immunohistochemistry for the met-enkephalin we found about half of biocytin filled neurons were enkephalin positive (8/15 neurons, Fig S3. B, C), consistent with described populations of enkephalinergic D2-receptor containing medium spiny neurons [25]. Reconstruction of biocytin-filled neurons revealed that enkephalinergic ASt MSN’s frequently extended dendrites towards the ASt-LA border with dendrites terminating at the border or projecting into the LA proper (4/15 neurons terminating in the LA, 6/15 terminating at the border, Fig. S3, B, C). Whilst some direct dendritic connectivity exists [36], the differences in amplitude and rise time of opioid signals favour diffusion rather than direct synaptic release as the primary mechanism for enkephalin delivery in the LA.

In summary, these findings indicate that auditory thalamic activity is sufficient to drive enkephalin release from D2-expressing MSNs in the ASt, which can diffuse to, and modulate signalling in the LA and thus potentially influence fear learning.

### Enkephalin release decays along a diffusion gradient across the ASt-LA network

A key question is whether enkephalin concentrations at different distances from the ASt release site are sufficient to engage opioid receptors. Previous work in the intercalated cells of the amygdala demonstrated that dense-core vesicles containing met-enkephalin are most commonly found apposed to axons and axon terminals [37], suggesting that enkephalin release occurs at somatodendritic sites where local concentration would be the highest. Because δLight1.1 contains a somatic restriction tag that restricts its expression to cell bodies [28], it is likely positioned away from these release sites and may therefore underestimate true release concentrations. We therefore performed parallel experiments using the µ-opioid receptor-based biosensor UMASS, which lacks the somatic restriction tag of δLight1.1 and is therefore expressed at higher levels on dendrites and axons than the cell body targeted δLight1.1 (Fig 4. A, B) [28, 38]. Because both sensors bind enkephalin with high affinity [26], the identity of the detected peptide is not altered by the choice of sensor.

**Figure 4.**
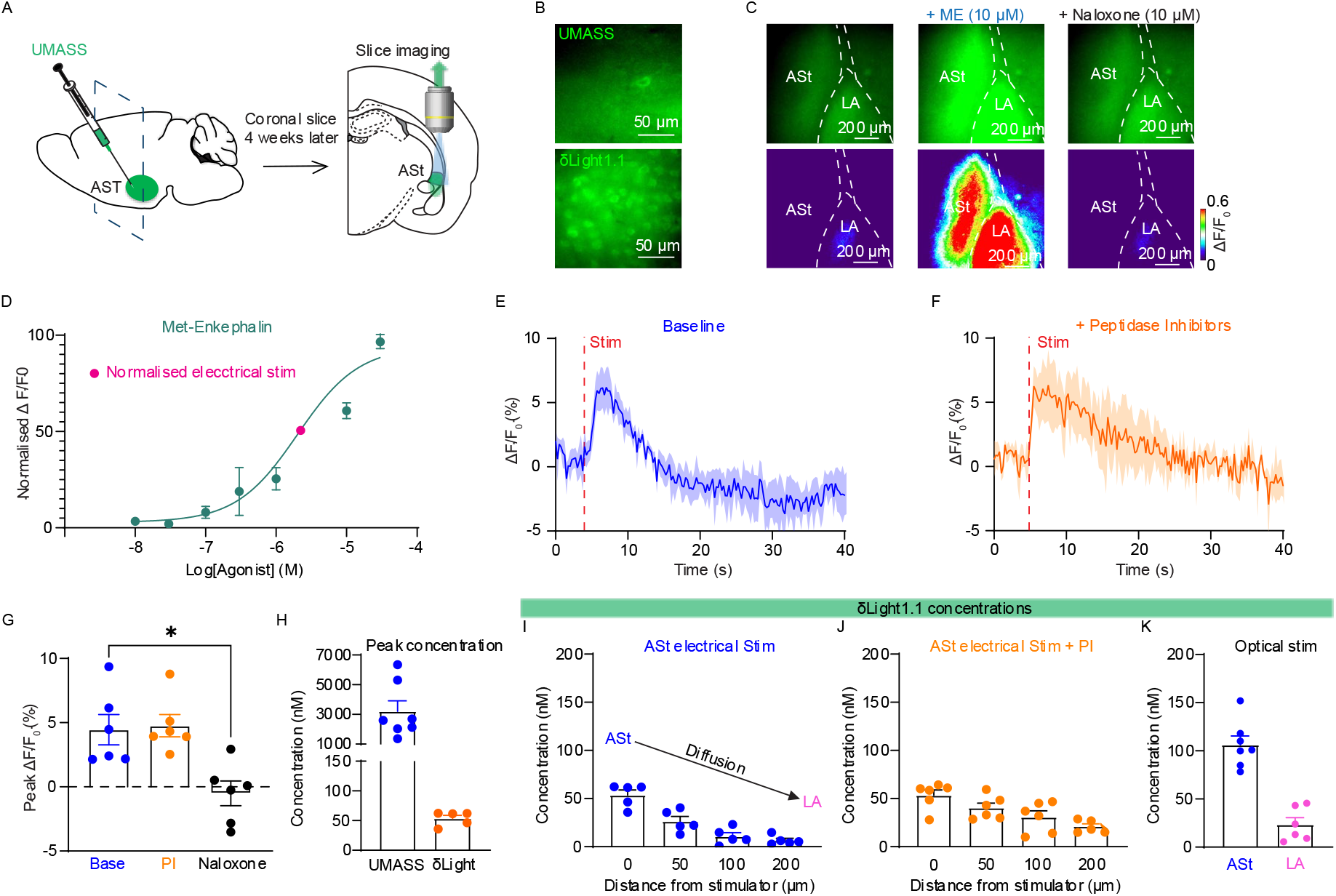
Enkephalin is released at a high concentration that diffuses down the ASt to the LA. **(A)** Schematic diagram showing injection of UMASS into the ASt and recording location in the ASt. **(B)** Representative pseudo-coloured images showing expression of UMASS (top) and δlight1.1 (bottom) in the ASt. Expression of UMASS can be seen on somatic processes whilst δlight1.1 expression is limited to the cell soma. **(C)** Pseudo-coloured UMASS images and heatmap showing increases in fluorescence following bath application of met-enkephalin (10 µM) and reversal by naloxone (10 µM) **(D)** Normalised UMASS ΔF/F_0_ concentration-response curves for met-enkephalin, normalised response for electrical stimulation shown in pink. **(E-F)** Average traces showing ΔF/F_0_ changes evoked by optical stimulation (100 stimulations at 50 Hz) as measured in the ASt during baseline and peptidase inhibitor. **(G)** Chart showing peak ΔF/F_0_ responses measured as an average of the first 8 frames post-stimulation in the ASt during baseline, peptidase inhibitor and naloxone application. (Two-way repeated measures ANOVA, Sidak’s multiple comparison, Baseline vs Naloxone: P = 0.05) **(H)** Chart showing fitted concentrations measured at stimulation site for UMASS and δLight1.1. **(I-K)** Charts showing fitted δLight1.1 concentrations measured at various distances, with and without peptidase inhibitors and for optical stimulation in the ASt and LA.

Similar to δ-light1.1 bath application of met-enkephalin (10 μM) caused an increase in UMASS fluorescence which was abolished by application of the antagonist naloxone (10 μM) (Fig 4, C). Stimulation-evoked UMASS signals were significantly larger than deltaLight1.1 signals (Electrical: 1.78 ± 0.21% peak ΔF/F0, n = 4; Optical: 3.74 ± 0.71% peak ΔF/F P = 0.025, Unpaired Student’s t-test) possibly reflecting UMASS’s closer proximity to release sites. When the UMASS signal was fitted onto the UMASS concentration response curve the estimated extracellular enkephalin concentration was ∼ 3 µM at the stimulation site (Fig 4. H). Whereas fitting stimulation-evoked δLight1.1 responses onto its met-enkephalin concentration response curve (Fig 2. F) yielded lower estimated enkephalin concentrations of ∼ 53 nM at the site of electrical stimulation, declining to ∼ 6.3 nM at distal sites measured in the LA (Fig 4, I). Peptidase inhibitor application did not increase peak concentrations but allowed the concentration of enkephalin to remain above 20 nM at distal sites (Fig 4. J). Optical MGN stimulation evoked approximately double the concentration of electrical stimulation (∼100 nM), with hotspots reaching ∼160 nM (Fig 4. K) and approximately 20 nM in the LA. The greater effectiveness of synaptically driven release over electrical stimulation is consistent with our previous findings in the intercalated cells [37], and suggests either that the strong synaptic drive excites more enkephalinergic MSNs or that glutamate acts through its range of receptors to trigger peptide release.

These calculated extracellular enkephalin concentrations are well above the K_i_ of met-enkephalin at sub-nanomolar μ-opioid receptors [39], indicating they will achieve substantial receptor occupancy and within range producing functional effects in the amygdala, as 100 nM met-enkephalin produces a half-maximal inhibition of glutamatergic transmission at the BLA-ITC synapse [37]. Together these data indicate that enkephalin is released at high local concentrations that decay rapidly with distance, that this decay is actively sculpted by peptidase activity, and that concentrations across the ASt-LA network remain sufficient for functional µ-opioid receptor engagement.

### Reducing enkephalin enhances fear learning

To directly test the contribution of enkephalin in the ASt during fear learning, we developed a viral CRISPR-based approach targeting the proenkephalin (PENK) gene in rats. A single-guide RNA targeting exon 2 of the rat proenkephalin gene was cloned and packaged into AAV-SaCas9 vectors [40]. Similarly, a non-targeting guide was also developed. Injections into the ASt reduced met-enkephalin immunoreactivity by approximately 25% compared to non-targeting controls (P < 0.0001, Fig 5 A-C), with the reduction extending across the rostral-caudal ASt without affecting met-enkephalin expression in the CeA (Fig S4 A).

**Figure 5.**
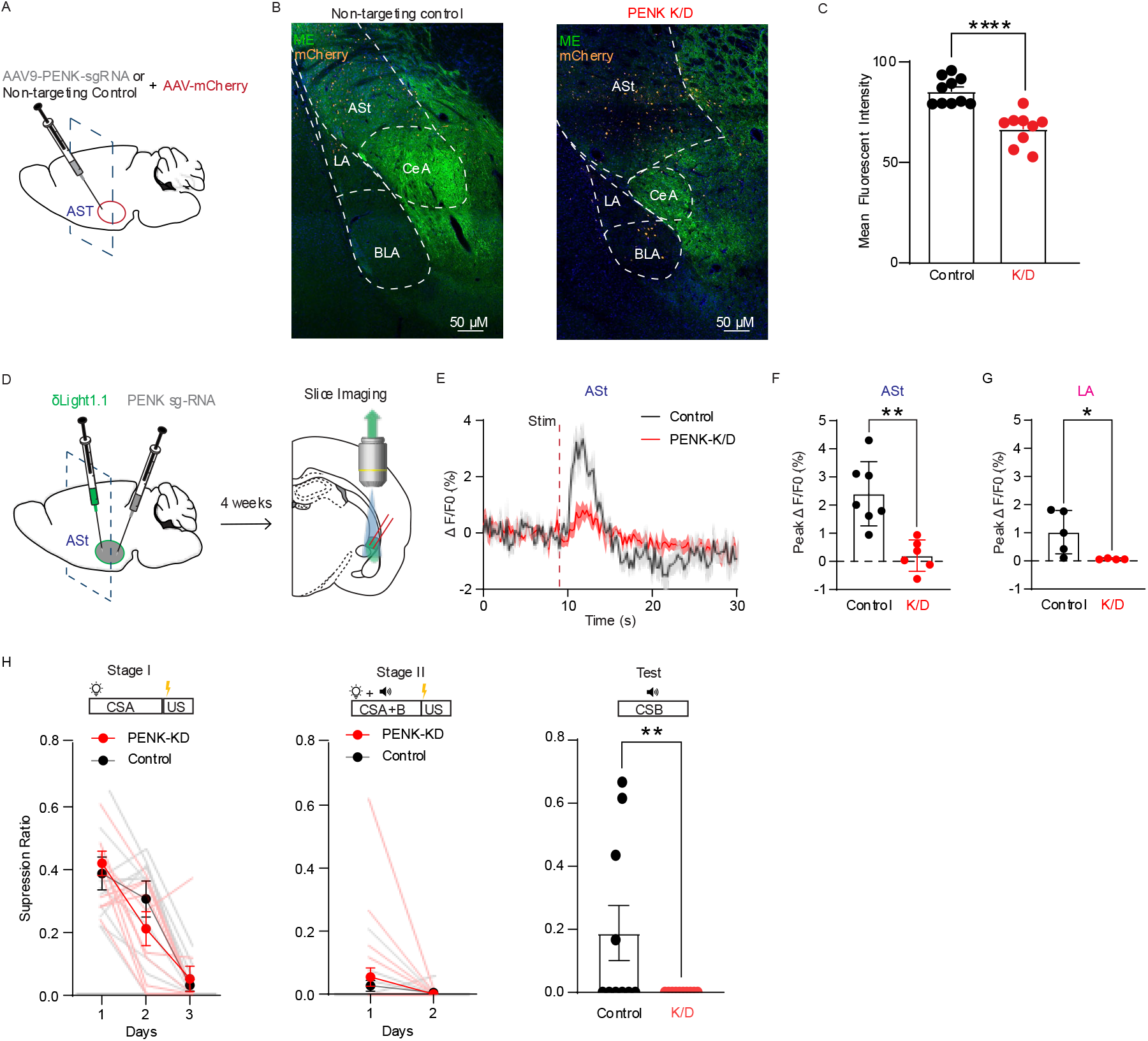
Enkephalin released from the ASt constrains fear learning. **(A**) Schematic diagram showing injection of AAV-PENk-sgRNA or non-targeting control and AAV-mCherry into the ASt. **(B)** *Post hoc* confocal images showing DAPI (blue) mCherry (red) and met-enkephalin (green) expression in the ASt amygdala regions for non-targeting controls injections and PENK-K/D. **(C)** Graph showing mean fluorescent intensity as measured in ASt slices taken from knockdown (*n = 9*) and control (*n = 10*) animals. **(D)** Schematic showing co-injection of δLight1.1 and AAV-PENK-sgRNA in the ASt. Slices were taken 4 weeks post-injection of fluorescent recordings. **(E)** Average traces showing ΔF/F_0_ changes evoked by local electrical stimulation (100 stimulations at 50 Hz) as measured in the ASt for control (black) and knockdown (red) slices. **(F-G)** Chart showing peak ΔF/F_0_ responses measured as an average of the first 8 frames post-stimulation in the ASt and LA during baseline for control and knockdown slices. (Two-tailed unpaired *t-*test, ASt: P = 0.001; LA: P = 0.04). Data presented as mean ± s.e.m with individual data points. **(H)** Suppression ratios during the associative blocking paradigm. In stage I, animals were conditioned to a single visual cue (CSA). In stage II, animals were conditioned to a compound stimulus consisting of both auditory (CSA) and visual cues (CSB). At test, animals were presented with the second stimulus (CSB) alone. Control animals showed robust blocking, with high suppression ratios to CSB, whereas PENK knockdown animals showed significantly lower suppression ratios to CSB, indicating enhanced fear learning to the added cue despite prior conditioning to CSA

To verify the functional impact of PENK reduction on endogenous opioid signalling we co-expressed the opioid sensor δLight with PENK knockdown virus and measured stimulation evoked fluorescence responses in acute brain slices (Fig. 5 D). In control non-targeting animals, electrical stimulation of the ASt reliably evoked changes in δLight signal in the ASt similar to response seen previously (Fig. 2 F). Changes in fluorescence evoked by ASt electrical stimulation were also seen in the LA but with a smaller amplitude consistent with a peptide signal declining as it diffuses across the ASt to the LA (Fig. 5 D). In slices from PENK knockdown animals, stimulation evoked responses were significantly attenuated in both the ASt and the LA. Estimated enkephalin concentrations following knockdown were 5 nM in the ASt and 0.7 nM in the LA, reflecting this decrease in signal. Critically, the complete abolition of LA opioid signals following knockdown targeted exclusively to the ASt indicates that the opioid signals seen in the LA originates from enkephalin released in the ASt rather than from local release within the LA itself. The reduction in δLight fluorescence scaled with knockdown efficiency across slices (R^2^ = 0.61, Fig. S4 B), indicating that signal attenuation reflected the degree of enkephalin depletion rather than an off-target viral effect. Together these data demonstrate that the PENK knockdown selectively reduced enkephalin availability in the ASt and that opioid signalling in the LA is dependent on enkephalin originating from this region.

The behavioural consequences of altered enkephalin signalling were examined using this viral PENK knockdown approach. If ASt enkephalin normally acts to limit fear learning, then reducing it should enhance the fear a cue can acquire. This is difficult to test directly, because fear learning rapidly approaches ceiling effects, so simple acquisitions can become insensitive to further enhancement. We therefore used a blocking-like design, which constrains learning to a target cue and so provides a low baseline against which enhanced learning can be detected. In Stage I, rats were fear conditioned to a single cue (CSA). In Stage II, the same rats were conditioned to a compound stimulus (CSA+B), in which CSA was presented together with a novel cue (CSB), the compound stimulus co-terminating with footstock. Rats were then tested with CSB alone. This design establishes conditions under which learning to CSB is normally constrained, allowing us to ask whether removing enkephalin restores it. In intact animals, prior conditioning to CSA limits the learning that CSB can acquire as since CSA already predicts the outcome, little prediction error remains to drive new learning about the added cue [41]. Consistent with this prediction, control rats injected with non-targeting virus showed robust blocking acquiring little fear to CSB (Fig. 5 H). In contrast, rats with ASt PENK knockdown showed significantly greater fear to CSB (two tailed unpaired *t*-test, P = 0.045, Fig. 5 H), indicating that reductions in ASt enkephalin enhanced learning about CSB, consistent with enkephalin normally limiting the strength of fear learning.

Together these data demonstrate that ASt-derived enkephalin supports opioid signalling across the ASt-LA circuit and that its removal is sufficient to enhance associative fear learning, establishing a causal link between enkephalin release from the ASt and the regulation of fear memory formation.

### Dopamine is released in the LA through volume transmission from the ASt during fear learning

The spread of enkephalin from the ASt into the LA positions it to regulate a range of targets within fear learning circuits. Dopamine is released in the LA during fear learning [42, 43], where it drives synaptic plasticity at sensory inputs and strengthens cue-threat associations [4], making dopamine a key candidate target of ASt enkephalin signalling.

To characterise dopamine dynamics in the LA during fear learning, we expressed GrabDA3mR in the BLA and recorded dopamine signals during auditory fear conditioning (Fig 6 A, B). Robust dopamine signals were detected in the LA during fear conditioning, with responses transferring form the US to the CS as training progressed (Fig 6, C) Compared to the δLight seen previously, dopamine signals lasted notably longer than 10 seconds. These data demonstrate that dopamine is dynamically recruited in the LA during fear learning, that its signalling, like that of enkephalin, tracks the predictive value of the auditory cue. This coincident release suggests that enkephalin could target dopamine release during fear learning.

**Figure 6.**
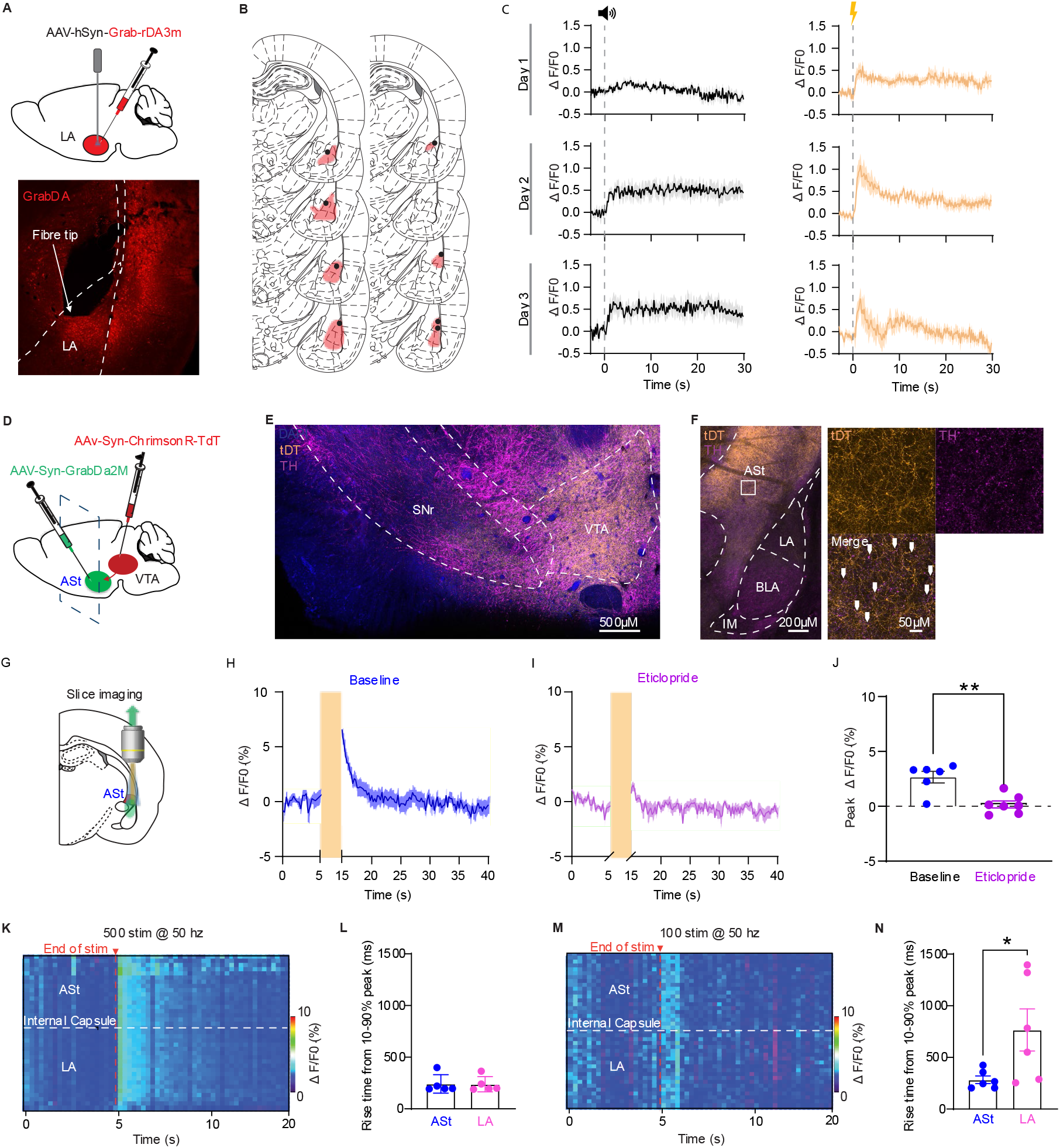
Dopamine is released in the LA during fear learning and spreads via volume transmission from the ASt. **(A)** Top: Schematic showing injection of AAV-GrabDA3mR into the LA and optical fibre tip implanted into the same region. Bottom: Representative image showing GrabDAr expression and fibre tip location in the LA. **(B)** Schematic diagrams of LA GrabDAr expression and fibre tip placements for all animals **(C)** Average GrabDAr traces during tone (black) and shock (yellow) across learning days. **(D)** Schematic showing injection of AAV-syn-GrabDA2m in the ASt and AAV-syn-ChrimsonR-tDt into the ventral tegmental area. **(E)** *Post hoc* images showing expression of DAPI (blue) ChrimsonR-tDT (orange) and tyrosine hydroxylase (magenta) in the midbrain. ChrimsonR cell bodies primarily located in the VTA. **(F)** *Post hoc* images showing tyrosine hydroxylase and ChrismonR-TdT terminal expression in the amygdala, with higher expression in the ASt relative to LA. High magnification of white box in the ASt shows co-expression of ChrimsonR-tDT and tyrosine hydroxylase. **(G)** Schematic showing imaging location in the ASt. **(H-I)** Average ΔF/F_0_ traces showing dopamine transients evoked by high intensity optical stimulation (500 pulses at 50 Hz and 1 ms pulse width), measured in the ASt during baseline and eticlopride (10 uM) application. **(J)** Chart showing peak ΔF/F_0_ responses measured as an average of the first 8 frames post-stimulation during baseline and eticlopride application. **(K)** Heatmap showing spatial-temporal profile of the evoked fluorescence in the ASt and LA during high intensity stimulation. **(L)** 10-90% 10-90% rise times measured in the ASt and LA. **(M-N)** Same as but for lower stimulation (100 pulses at 50 Hz and 1 ms pulse width).

Anatomical tracing using ChrimsonR injection into the VTA revealed dense dopaminergic innervation of the ASt confirmed by tyrosine hydroxylase and tdTomato co-labelling (Fig 6. D-F), but considerably sparser innervation of the LA, consistent with previous tracing studies [11, 23]. Whole cell recording revealed sparse fast excitatory connectivity (1/15 directly connected, Fig S5, A-C), similar to connectivity rates seen previously [44]. The sparse LA innervation raised the question of whether dopamine signals in the LA arise entirely from local VTA terminals or whether ASt-released dopamine contributes via volume transmission. To determine this, we expressed ChrimsonR in VTA neurons and the dopamine sensor GrabDA2H in the ASt to monitor changes in extracellular dopamine evoked by VTA stimulation (Fig 6D). Optical stimulation of VTA terminals evoked robust eticlopride sensitive dopamine transient (Fig 6. G-J). Unlike enkephalin, dopamine transients in the LA exhibited no onset delay or difference in response size relative to the signals observed in the ASt (Fig 6 K, L). However, given the high stimulation that was used in this experiment, we reasoned that this stimulation could be occluding the initiation kinetics of the dopamine response. Therefore, we repeated the experiment using a shorter stimulation (100 stimulations at 50 Hz). Under these conditions, LA dopamine signals exhibited a significantly slower rise time compared to those in the ASt (Fig 6 M, N). Together with the sparse VTA innervation of the LA relative to the ASt, these data indicate that a component of the LA dopamine signal likely reflects diffusion from the ASt via volume transmission, potentially extending the neuromodulator hub function of the ASt beyond enkephalin to encompass dopamine itself.

### Enkephalin supresses dopamine release in the ASt through activation of μ-opioid receptors

Both enkephalin and dopamine are released within the ASt during fear learning and appear capable of reaching the LA via volume transmission, raising the possibility that these two neuromodulator systems interact within this shared extracellular space. Such an interaction would provide a mechanistic link between opioid release and its established role in constraining fear learning [11, 43]. Using the combined GrabDA2h and ChrimsonR expression system we used in our previous experiments, we tested whether opioid receptor activation modulates dopamine release from these terminals. First, bath application of met-enkephalin (10 µM) strongly suppressed stimulation-evoked dopamine release (Fig 7 A-C). This inhibition was fully reversed after application of the selective µ-opioid receptor antagonist CTAP (Fig 7 A-C), indicating that enkephalin suppresses dopamine release in the ASt via µ-opioid receptor activation. In contrast, application of the selective κ-opioid receptor agonist U69 (3 μM) had no effect on stimulation-evoked dopamine signal, and nor did the ϰ-opioid receptor antagonist norBNI (10 nM), indicating that dopamine release from the VTA terminals in the ASt is insensitive to κ-opioid receptor activation. This differential sensitivity is consistent with previous work showing that BLA- and striatum-projecting VTA neurons represent distinct subpopulations with different opioid receptor profiles [24], prompting us to directly characterise ASt-projecting VTA neurons.

**Figure 7.**
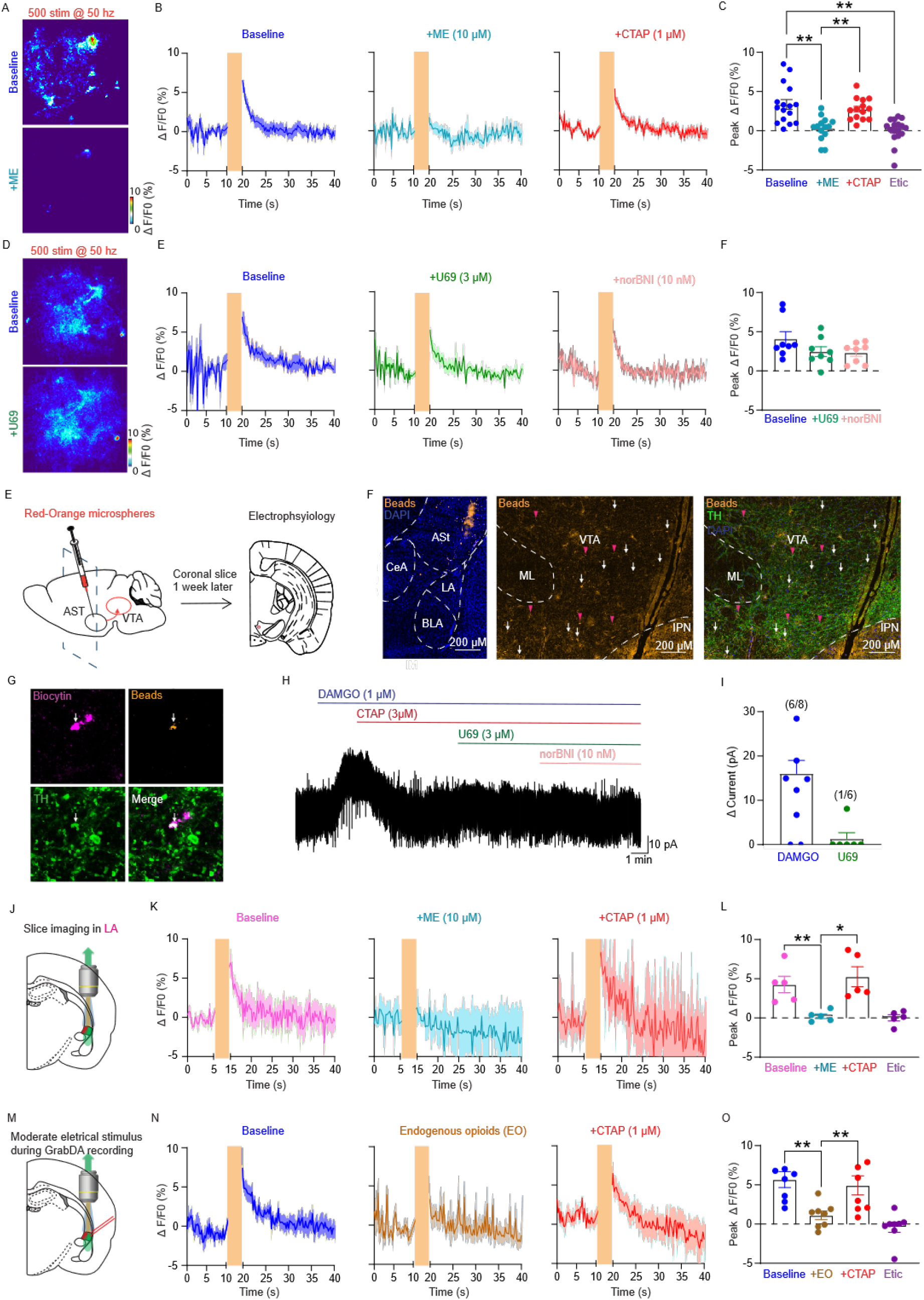
Enkephalin supresses dopamine release in then ASt and LA via activation of MOR. **(A**) Pseudo-coloured heatmaps showing ΔF/F_0_ following optical stimulation of VTA terminals during baselines (top) and met-enkephalin application (bottom) **(B)** Averaged traces showing ΔF/F_0_ during baseline, met-enkephalin and CTAP application. **(C)** Chart showing peak ΔF/F_0_ responses measured as an average of the first 8 frames post-stimulation during baseline, met-enkephalin (10 µM), CTAP (1 µM) eticlopride (10 µM) application. ** P < 0.01, One-Way ANOVA, Sidak’s multiple comparisons). **(D-F)** Same as (A-C) but for baseline, U69 (3 µM) and norBNI (10 nM). Schematic diagram showing injection of red-orange retrograde microspheres into the ASt. Slices were taken 1 week later for electrophysiology. **(E)** *Post hoc* images of the amygdala showing Red-orange microsphere injection in the ASt and microsphere expression localised in the VTA. White arrows indicate a bead positive ASt projecting neuron. Pink arrowheads show bead positive neurons that are co-localised with TH **(F)** High powered image of a patched neuron showing co-expression of biocytin, beads and TH. **(G)** Example trace showing outward potassium currents evoked by the selective µ-opioid receptor agonist DAMGO (1 µM) and the selective κ-opioid receptor agonist U69 in bead-positive VTA neuron from (F). **(H)** Chart showing change in current evoked by either DAMGO or U69 and the proportion of neurons sensitive to each agonist. **(I)** Schematic diagram showing imaging location in the LA **(J)** Averaged traces showing ΔF/F_0_ during baseline, met-enkephalin and CTAP application **(K)** Chart showing peak ΔF/F_0_ responses measured as an average of the first 8 frames post-stimulation during baseline, met-enkephalin, CTAP, eticlopride application. (*P < 0.05, ** P < 0.01, One-Way ANOVA, Sidak’s multiple comparisons) **(L-N)** Same as (J-K) but measured in the ASt with a moderate local electrical stimulation (5 pulses 150 Hz) to induce endogenous opioid released combined with high intensity optical stimulation of VTA terminals.

**Figure 8.**
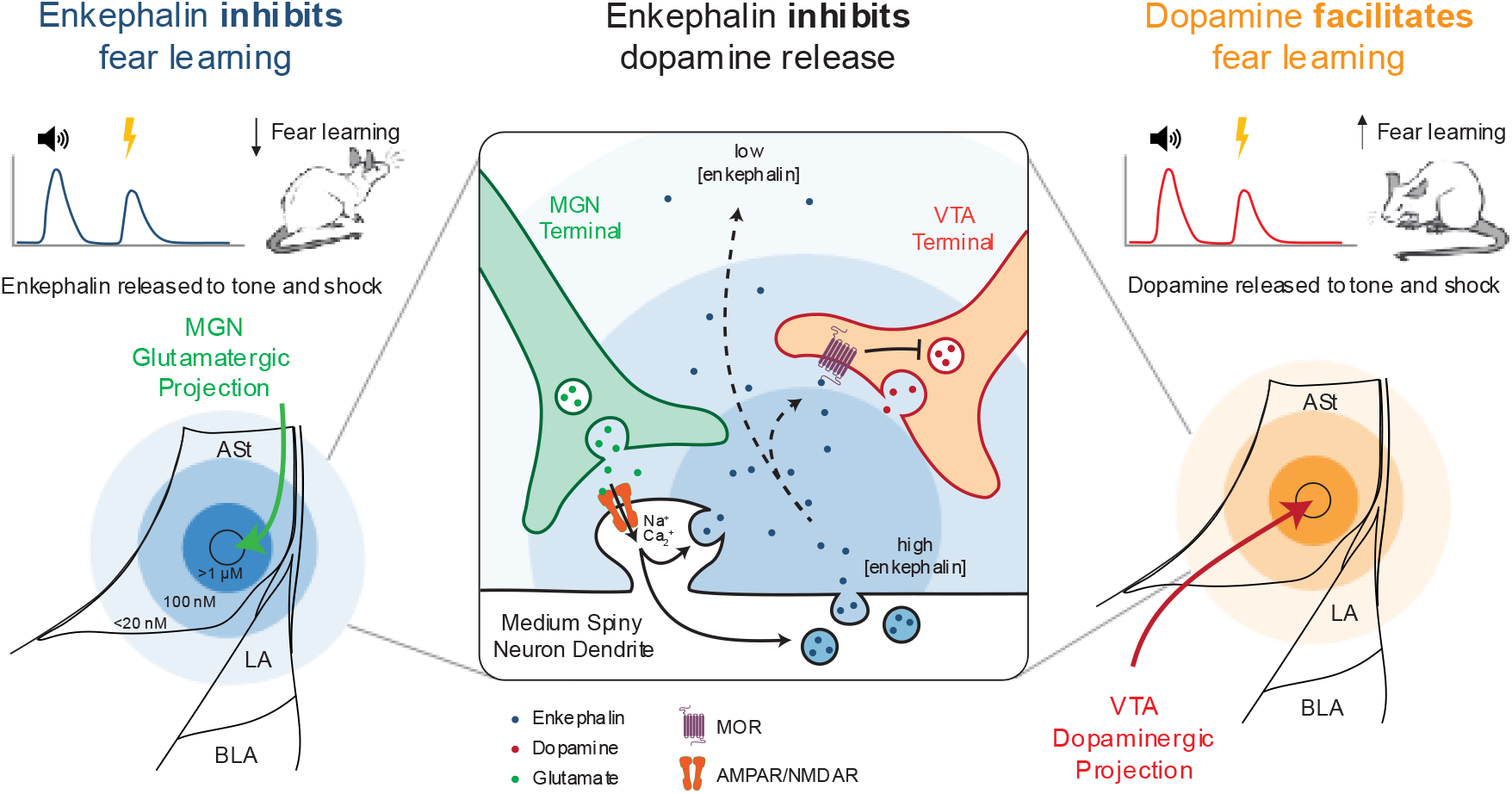
Proposed model

To address this directly, we injected fluorescent retrograde microspheres into the ASt and performed whole-cell patch clamp recording from bead-positive neurons in the VTA and determine their opioid sensitivity (Fig 7 D, E). The µ-selective agonist DAMGO evoked outward K^+^ currents in 6/8 neurons (Fig 7, F-H), whereas application of a selective κ-opioid receptor agonist U69 evoked outward K^+^ currents in only 1/6 bead-positive neurons (Fig 7, F-H). A proportion of bead positive neurons sensitive to DAMGO were also tyrosine hydroxylase positive (Fig 7 F), indicating that ASt-projecting dopaminergic neurons are predominantly µ- sensitive, consistent with the BLA-projecting rather than striatum-projecting population [24]. Importantly, whilst opioid inhibition of striatal dopamine occurs indirectly via µ-opioid receptors [24], our data demonstrate direct presynaptic µ-opioid receptor mediated inhibition of dopamine terminals in the ASt, representing a more direct mechanism of opioid control over dopamine release.

Since enkephalin released from the ASt diffuses into the LA, we next asked whether this signal can suppress dopamine release within the LA itself. Bath application of met-enkephalin (10 µM) strongly inhibited stimulation-evoked dopamine release in the LA, fully reversed by CTAP (Fig 7 I-K), demonstrating the same µ-opioid receptor mechanism identified in the ASt.

Finally, to test whether endogenous enkephalin released from the ASt is sufficient to regulate dopamine signalling, we electrically stimulated (5 pulses at 150 Hz) the ASt whilst optically stimulating VTA terminals (Fig 7 L). In the presence of peptidase inhibitors (10 μM Bestatin, 10 μM Thiorphan and 1 μM Captopril), ASt stimulation significantly reduced VTA terminal driven dopamine transients (Fig 7 L-N). This suppression was fully reversed by CTAP (Baseline: 5.67±1.01% ΔF/F_0_; with ASt stimulation: 1.11±0.56% ΔF/F_0_; P = 0.04; with CTAP: 4.93±1.22% ΔF/F_0_; P = 0.04; *n* = 8, Fig 7 L-N). Thus, endogenously released enkephalin from the ASt is sufficient to supress dopamine release via µ-opioid receptors.

Together these data demonstrate that ASt-derived enkephalin, acting via volume transmission, suppresses dopamine release across both the ASt and LA through µ-opioid receptor activation, providing a mechanism through which the ASt can broadly regulate the dopaminergic signals that drive fear learning.

## Discussion

A fundamental but previously unresolved questions is how endogenous opioids regulate aversive learning. Here we show for the first time that enkephalin is dynamically released within the amygdala during fear conditioning, originating from enkephalinergic medium spiny neurons of the ASt, and spreading via volume transmission across the ASt-LA border to limit dopamine release and constrain fear learning. These findings position the ASt not as a passive transition zone between amygdala and striatum, but as a functionally distinct neuromodulator hub.

A central finding of this study is the robust recruitment of the ASt by the MGN. Whilst the MGN is classically viewed as a sensory relay to the LA [8, 33, 34], our data reveal that MGN input engages enkephalinergic ASt neurons more strongly than LA neurons. The MGN undergoes associative plasticity during fear learning, with CS-evoked responses strengthening across conditioning [45], providing a mechanism through which increasingly strong thalamic drive progressively recruits enkephalin release as learning progresses and the opioid signal shifts from the US to the CS.

The nature of neuropeptide signalling has been proposed to be fundamentally different to the point-to-point communication of neurotransmitters, such as glutamate, with neuropeptides potentially relying on volume transmission and having a wider sphere of influence. Several lines of evidence from this study support this proposal with ASt-derived enkephalin acting via volume transmission in the LA. Firstly, MGN-driven opioid signals peaked more quickly in the AST than the LA. Secondly, the MGN-driven opioid signals were larger in the AST then decreased down a concentration gradient into the LA, consistent with the highest concentration at the AST release site. These characteristics are consistent with the concentration dissipation and movement of enkephalin described using voltammetry for endogenous enkephalin [46], and electrophysiological responses to photoactivatable leu-enkephalin, where enkephalin moved 200 µm within a second [47]. Finally, and most directly, knocking down enkephalin expression in the AST nearly abolished LA opioid signals, providing causal evidence that the LA signal originates from ASt enkephalin release rather than local LA sources. Together this data demonstrates that enkephalin is released and through volume transmission influences local cells and neighbouring brain regions.

Knock down of enkephalin in the LA was sufficient to enhance fear learning in a blocking paradigm demonstrating that ASt-derived enkephalin normally exerts a meaningful constraint on the strength of associative fear memories. Classically, endogenous opioids have been viewed primarily in the context of pain modulation during aversive learning [17]. Our data support a broader view, whereby enkephalin release from the ASt functions as a negative learning signal that limits the rate of fear memory formation within the LA without preventing it entirely.

Dopamine in the LA is a key downstream target of this enkephalin signal. Enkephalin acting at µ-opioid receptors directly suppresses dopamine release in both the ASt and LA and endogenously released enkephalin from ASt stimulation is sufficient to produce this suppression. Importantly, this represents a more direct mechanism of opioid control over VTA driven dopamine release than in striatal circuits, where µ-opioid receptor activation does nor inhibit VTA driven dopamine release but only inhibits dopamine release under specific conditions when it is driven by cholinergic neurons [48]. The ASt-projecting VTA population shares the same µ-sensitive and largely κ-insensitive profile of the amygdala-projecting rather than striatum-projecting neurons [24], meaning enkephalin selectively suppresses dopamine release in the amygdala fear circuits. By dampening dopamine release during conditioned or unconditioned stimulus presentation, ASt derived enkephalin may reduce the reinforcement signal that drives fear memory consolidation in the lateral amygdala [49], consistent with the enhancement of fear learning following enkephalin knockdown. Furthermore, if clinical µ-opioid agonists such as morphine and fentanyl engage the same presynaptic mechanism, providing a direct route through which opioid drug use could alter fear circuit dopamine signalling.

The finding that both enkephalin and dopamine may operate via volume transmission repositions the ASt as a neuromodulator broadcasting hub whose influence is governed by release, diffusion and degradation rather than synaptic connectivity. The concentration gradients established by volume transmission may themselves carry functional information, has been recently shown in dopaminergic signalling, high local concentrations engage membrane-delimited Gβγ signalling whilst lower distal concentrations engage cAMP-dependent pathways [50]. Pathological reduction in enkephalinergic tone, such as those produced by opioid withdrawal-driven upregulation of peptidase activity [35] or chronic pain-induced suppression of opioid release [51], would not simply reduce signal magnitude but shift the balance between distinct intracellular signalling cascades across the ASt-LA network. Additionally, whilst the present study focused on auditory fear learning, the ASt receives convergent input from visual, somatosensory, gustatory, olfactory and limbic structures [52], suggesting it may represent a general mechanism through which diverse sensory experience engage enkephalin release to limit associative learning

In summary, these findings identify the ASt as an active regulator of fear memory operating through thalamic sensory integration, enkephalin volume transmission and dopaminergic modulation. Endogenous opioids act not merely as modulators of affect but as learning signals that constrain the strength of aversive associations, with the ASt setting the gain of fear memory formation. Understanding how this circuit is disrupted under conditions of chronic stress or opioid exposure will be important for addressing the dysregulation of fear learning in psychiatric disease.

## Methods

### Animals

Male Sprague-Dawley rats (4-12 weeks old) and male Long-Evans rats (10-12 weeks old) were obtained from the Animal Resources Centre (Ozgene ARC, Australia). Animals were housed in groups of up to six, maintained on a 12/12-hour light-dark cycle, and provided with food and water ad libitum. All experiments were performed under protocols approved by the University of Sydney Animal Ethics Committee (1637/2019) and the University of New South Wales, Sydney.

### Viral constructs

AAV8-Syn-ChR2(H134R)-GFP (Addgene plasmid #58880), AAV9-hSyn-GRAB-DA2h (Addgene plasmid #140554), AAV2/9-hSyn-GRAB-DA3mR (Addgene plasmid #208702) and AAV5-Syn-ChrimsonR-tdT (Addgene plasmid #59171) were obtained from Addgene. AAV9-hSyn-δLight1.1 was a gift from Lin Tian. AAV5-hsyn_HA-Flas-uMASS_1_WRPE (Addgene plasmid #209760) was a gift from Michael Bruchas. AAV9-PENK-KO-sgRNA1 (5’- GAAACCGCCATACCTCTTGGC-3’) and AAV9-control non-targeting sgRNA (5’- GGAGACGAATAATGCGTCTCC-3’) vectors were generated in-house, validated by sequencing and packaged by the Children’s Medical Research Institute (Western Sydney Local Health District, Australia). Fluorescent retrograde microspheres (FluoSpheres, red/orange, catalog#F8794) were obtained from Thermofisher.

### Stereotaxic injections

Rats aged 4-6 weeks were anaesthetised with 5% isoflurane for induction and maintained at 1.5-3% isoflurane mixed with oxygen (1 L/min) throughout surgery. Animals received subcutaneous carprofen (5 mg/kg) and bupivacaine (5%) before a midline incision was made and craniotomies performed above the target regions. Injections were delivered through pulled glass pipettes at the following coordinates relative to bregma (anteroposterior, mediolateral, dorsoventral, in mm): lateral amygdala (- 2.5, ± 5.2, - 7.4), amygdalo-striatal transition zone (- 2.4, ± 4.6, - 7.4), medial geniculate nucleus (- 5.4, ± 3.1, - 6.2), and ventral tegmental area (- 5.3, ± 0.8, - 8.2). Pipettes were left in place for 5 minutes after injection before retraction. Craniotomies were sealed with bone wax (Coherent Scientific, AUS) and incisions closed with silk sutures. Animals received intraperitoneal cephazolin (100 mg/ml, Hospira, AUS) and 1 ml normal saline and were allowed to recover for at least 7 days before further procedures.

### Brain slice preparation

Rats were deeply anaesthetised with isoflurane, decapitated, and the brain rapidly removed into ice-cold cutting artificial cerebral spinal-cord fluid (aCSF) containing (in mM): 125 NaCl, 25 NaHCO3, 11 D-glucose, 2.5 KCl, 1.25 NaH2PO4, 2.5 MgCl2, 0.5 CaCl2, continuously saturated with carbogen (95% O2, 5% CO2; 295 mOsm/L, pH 7.3). Coronal slices containing the amygdala (280 µm) were cut using a Leica VT1200 S vibratome and incubated in cutting solution at 34°C for 1 hour before recording.

### Electrophysiology

Slices were superfused with recording aCSF containing (in mM): 125 NaCl, 2.5 KCl, 1.25 NaH2PO4, 1 MgCl2, 2 CaCl2, 25 NaHCO3, 11 glucose, saturated with carbogen and maintained at 32-35°C. Neurons were visualised using an Olympus BX51WI upright microscope with differential interference contrast optics. Whole-cell recordings were made using patch pipettes (2-5 MΩ) filled with caesium chloride internal solution (in mM: 140 CsCl, 5 HEPES, 10 EGTA, 2 CaCl2, 2 Mg2ATP, 0.3 NaGTP, 3 QX-314.Cl, 0.05% biocytin) or potassium gluconate internal solution (in mM: 135 potassium gluconate, 8 NaCl, 2 Mg2ATP, 0.3 NaGTP, 11 EGTA, 10 HEPES, 0.05% biocytin), both at pH 7.3 and 280-285 mOsm/L. Series resistance was continuously monitored and experiments with changes greater than 20% were excluded. For optically evoked EPSCs, ChR2-expressing MGN terminals were stimulated with a single or paired 1 ms light pulse and recordings were made in the presence of picrotoxin and SR95531 to isolate glutamatergic currents. For retrograde bead-labelling experiments, opioid receptor agonist-evoked currents were assessed as outward potassium currents recorded at a holding potential of-60 mV. Signals were amplified using a Multiclamp 700B amplifier (Molecular Devices, USA), low pass filtered at 5 kHz, digitised at 10 kHz, and acquired using Axograph software (v18.0.0). All statistical analyses were performed using GraphPad Prism (v8.0.1), with results expressed as mean ± SEM and significance set at P < 0.05.

### Fluorescent slice imaging

Imaging was performed 4-6 weeks after injection of AAV9-hSyn-deltaLight or AAV9-hSyn-GRAB-DA2h. Slices were perfused with recording aCSF at 2.5 mL/min in an imaging chamber on an Olympus BX51WI microscope equipped with an Andor iXon Life 888 EMCCD camera (Oxford instruments, UK), controlled by Micro-Manager (NIH). Imaging used a x10/0.3 NA or x40/0.8 NA water-immersion objective. Green fluorescence was excited using a 475-495 nm filter and collected through a 510-540 nm emission filter; red fluorescence was excited using a 510-538 nm filter and collected through a 611-645 nm emission filter (Semrock, USA). Full-frame images (512 x 512 pixels) were captured at 4 Hz. For electrical stimulation, a bipolar electrode was positioned in the ASt. For optical stimulation, a x40 objective was positioned above the region of interest. Imaging and stimulation were synchronised using an Arduino board with custom software. Drugs were applied via bath perfusion at 2.5 mL/min. Images were analysed in FIJI by manually defining regions of interest and measuring mean grey value across frames. ΔF/F_0_ was calculated as (F_i_ −F_baseline_)/F_baseline_, where F_i_ is the fluorescence at each frame and F_baseline_ is the average fluorescence across the 20 frames preceding stimulation or 2 minutes preceding drug application. Spatial-temporal analysis was performed using custom-written software.

### Drugs

All stock solutions were prepared in Milli-Q water, except thiorphan, which was dissolved in DMSO (final working concentration of 0.01% DMSO). CNQX, DAMGO, CTAP, U69593, norBNI, deltorphin II, ICI 174,864 and D-AP5 were obtained from Tocris (Bristol, UK). Picrotoxin, SR95531, thiorphan, captopril, bestatin and methionine-enkephalin were obtained from Sigma-Aldrich (Missouri, USA). Dopamine was obtained from Cayman Chemicals (Michigan, USA). Eticlopride was obtained from HelloBio (Bristol, UK).

### Immunohistochemistry

Animals were deeply anaesthetised with isoflurane and given a lethal dose of intraperitoneal pentobarbital before transcardial perfusion with heparinised saline followed by 4% paraformaldehyde (PFA) in 0.1 M PBS. Brains were post-fixed overnight in 4% PFA at 4°C, washed in PBS, embedded in OCT, and stored at-80°C. Coronal sections (40 µm) were cut on a cryostat and stored in cryoprotectant at-30°C until processing. Sections were blocked in 10% normal goat serum, 0.5% BSA and 0.3% Triton X-100 in PBS, then incubated with primary antibodies: rabbit anti-GFP (1:1000, Invitrogen, catalog#A6455) for GFP-expressing tissue; rabbit anti-met-enkephalin (1:1000, Immunostar, catalog#20065) and sheep anti-tyrosine hydroxylase (1:1000, Millipore, catalog#AB1542) for peptide and dopaminergic terminal labelling. Sections were incubated with corresponding Alexa Fluor-conjugated secondary antibodies (donkey anti-rabbit 488 catalog#AB150073 or 568 catalog#AB175692; donkey anti-sheep 568 catalog#AB175712; all 1:500) alongside DAPI (1:1000), then mounted with Prolong Gold Antifade and imaged on a Nikon C2 confocal or Leica Thunder wide-field microscope (Sydney Microscopy and Microanalysis).

### Behavioural apparatus

All behavioural training was conducted in identical Med Associates chamber measuring 24 cm (length) x 30 cm (width) x 21 cm (height). Chambers were enclosed in a ventilated, sound-attenuating cabinets measuring 59.5 cm (length) x 59 cm (width) x 48 cm (height). A 3 W house light was mounted to the top right of the sidewall next to a speaker used to deliver auditory conditioned stimuli (CS). The left sidewall was fitted with a magazine dish where pellets (Bio-Serv) were delivered when a retractable lever located 4 cm to the right of the magazine was pressed. A metal grid was fitted to the floor of the chamber to deliver scrambled footshock as the unconditioned stimulus (US)

### Pavlovian fear conditioning

Male Long-Evans rats were food restricted to ∼85% of their body weight 2 days prior to behavioural training. This feeding regime was maintained until the end of the experiment. Two separate training contexts were used each day. In one context (reward), a lever was available to earn a food pellet reward delivered into a magazine. In the second context (neutral), a slate was slotted into the wall to block access to the magazine and level. The two context were further distinguished by different olfactory (peppermint and rosewater), auditory (fan on or off), visual (houselights on or off), as well as spatial (identical chambers in different locations) cues which were counterbalanced.

For pavlovian fear conditioning, rats first received daily lever press training in the reward context on Days 1-5 and were places in the neutral context for the same duration as training. Context order was randomised each day. On Days 1-2, rats would receive magazine training in the reward context, in these sessions reward was delivered for every lever press, with additional “free” reward delivery on a FI300 schedule. The free reward delivery facilitated learning of the lever press task. These sessions would terminate when the rat reached 100 lever presses or if 60 min had elapsed. On Day 3, rats lever pressed for reward on a VI30 schedule for 60 min. In all remaining sessions in the reward context, the rats lever pressed for the reward on a VI120 schedule in a 60 min session. On Days 9-10, rats were pre-exposed to auditory CS in the neutral context and a different auditory CS in the reward context. Auditory CSs were either a 60 s 10 Hz clicker or 60 s tone counterbalanced across the different context and rats. In each of these sessions, the 60 s CS was presented a total of 4 times on an intertrial interval (ITI) of 600-900 s per each 60 min. Rats were also tethered to the patch cable on these days. On Days 11-14, rats were tethered to patch cables for photometry recordings and then underwent fear conditioning in both contexts. For each day, the CS was presented four times and all presentation co-terminated with a 0.5 mA, 0.5 s footshock US. On Day 15, rats were tested for fear responses with four presentations of the CS alone.

### Blocking task

Rats underwent the same level training schedule as the pavlovian fear conditioning training for Days 1-4. Following 5 days of VI60 training, on Days 9-10 rats received pre-exposure to the CS. This consisted of 4 presentation of each CS in a randomized order across a 70 min session. CSA was a flashing LED and CSB was a clicker (85dB, 1Hz), with each CS presentation lasting 60 s. On Days 11-13, rats received Stage I training, where CSA co-terminated with a 0.5 s, 0.6 mA footshock. During each training sessions, CSA was presented 4 times. On Days 11-13, rats received Stage II training. In Stage II training, the CSA and CSB were presented simultaneously for 60 s and both co-terminated with the footshock. Both stimuli were presented together 4 times each session. On Day 16, rats were tested for fear responses to CSB alone, during the test, rats received 4 presentation of CSB alone without footshock. Fear responding was measured as suppression of lever pressing.

### Fibre photometry

All fibre photometry recordings were made using Doric lenses and Tucker-Davis Technologies systems (RZ5P, Synapse). For green δLight1.1 recordings a 465 and 405 nm wavelength light was used, emitted from LEDs controlled by programmable LED drivers. For red GrabDA3mR recordings an additional 560 nm wavelength light was used. Excitation lights were channelled through patch cables (0.39NA, ⌀400 mm core multimode prebleached patch cables). Light intensity at the tip of the cable was maintained at 10-13 μW across sessions. The experimental and isosbestic fluorescence signals were amplified and measured by Doric Fluorescence Detectors. Synapse software was used to control and modulate excitation lights (560 nm, 537 Hz; 495 nm, 209 Hz; 405 nm, 331 Hz). Signals were demodulated and low-pass filtered (3 Hz) in real time via the RZ5P. Behavioural events were recorded and sent to Med-PC via RZ5P/Synapse.

## Notes

### Competing Interest Statement

The authors have declared no competing interest.

### Summary of Updates

Updated figure 1 Panel A to reflect LA rather than BLA

### Funding

- National Health and Medical Research Council: APP1188908, APP1188949, Ideas GNT:2036815°
- Australian Research Council: DP250100345

