## Supplementary for "Enkephalin constrains fear learning via volume transmission to the lateral amygdala"

**Supplementary data**

**Supplementary Figure 1. Enkephalin is release from D2-expressing medium spiny neurons in the ASt.**

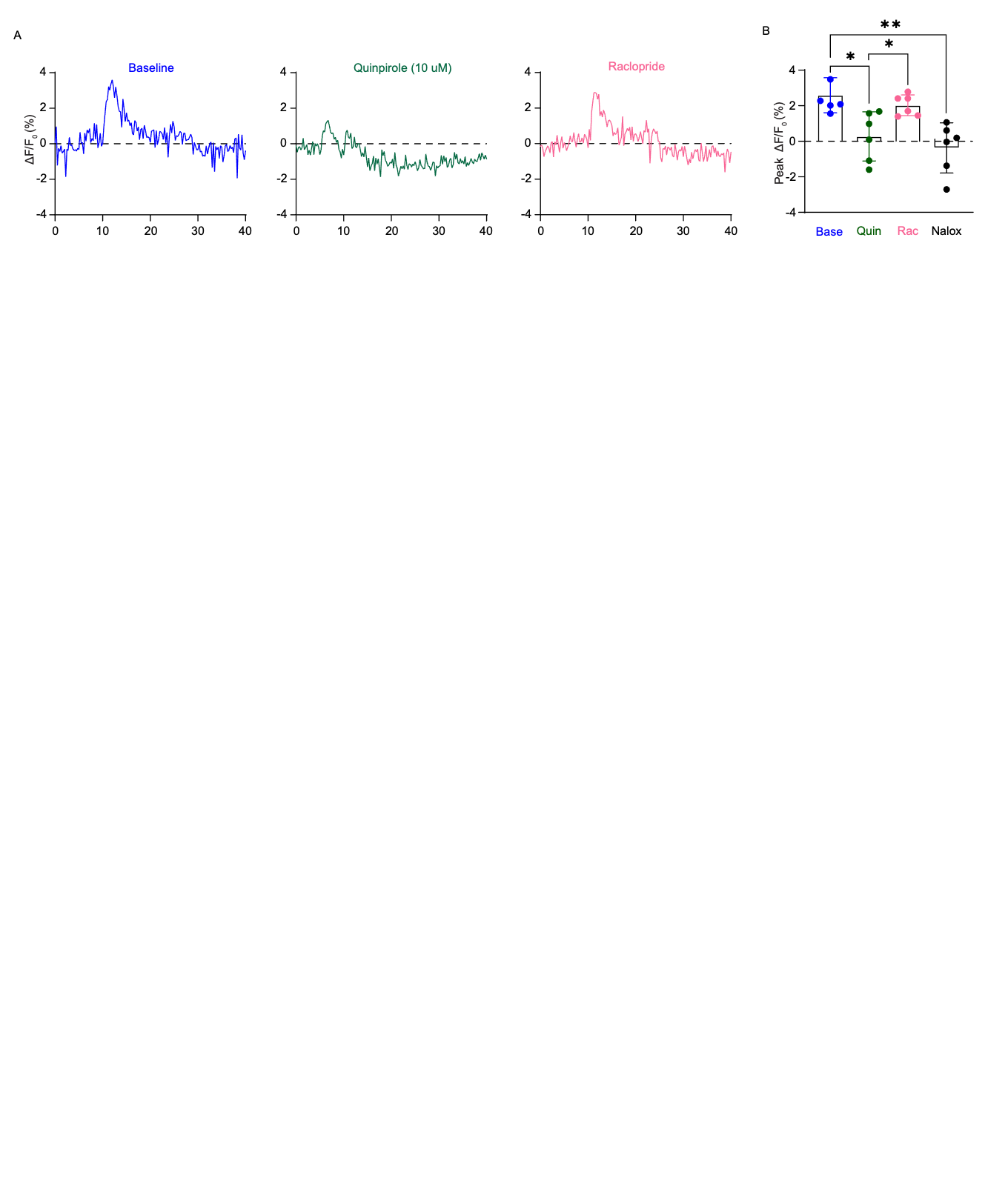

**(A)** Representative traces showing δLight1.1 ΔF/F_0_ during baseline, quinpirole and raclopride application. **(B)** Chart showing peak ΔF/F_0_ responses measured as an average of the first 8 frames post-stimulation during baseline and drug application. (*P < 0.05, ** P < 0.01, One-Way ANOVA, Sidak’s multiple comparisons)

**Supplementary Figure 2. Injection sites for ChR2-GFP and ChrimsonR-TdT**
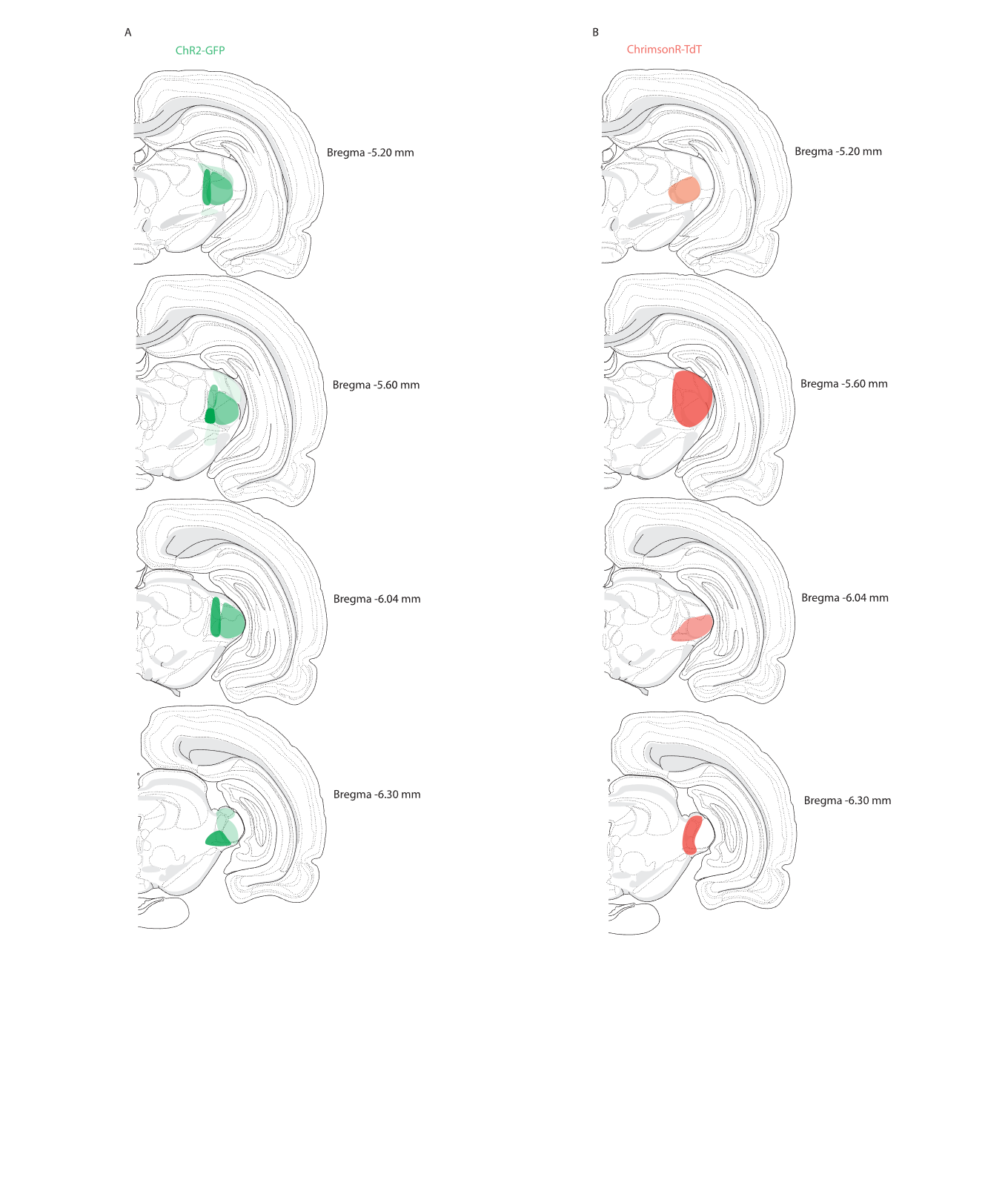

**(A-B)** Schematic diagram showing injection sites of injection at various rostral-caudal slices for ChR2-GFP and ChrimsonR-TDT. Dark colours indicate regions of high cell body expression.

**Supplementary Figure 3. ASt enkephalinergic sends processes towards the LA**

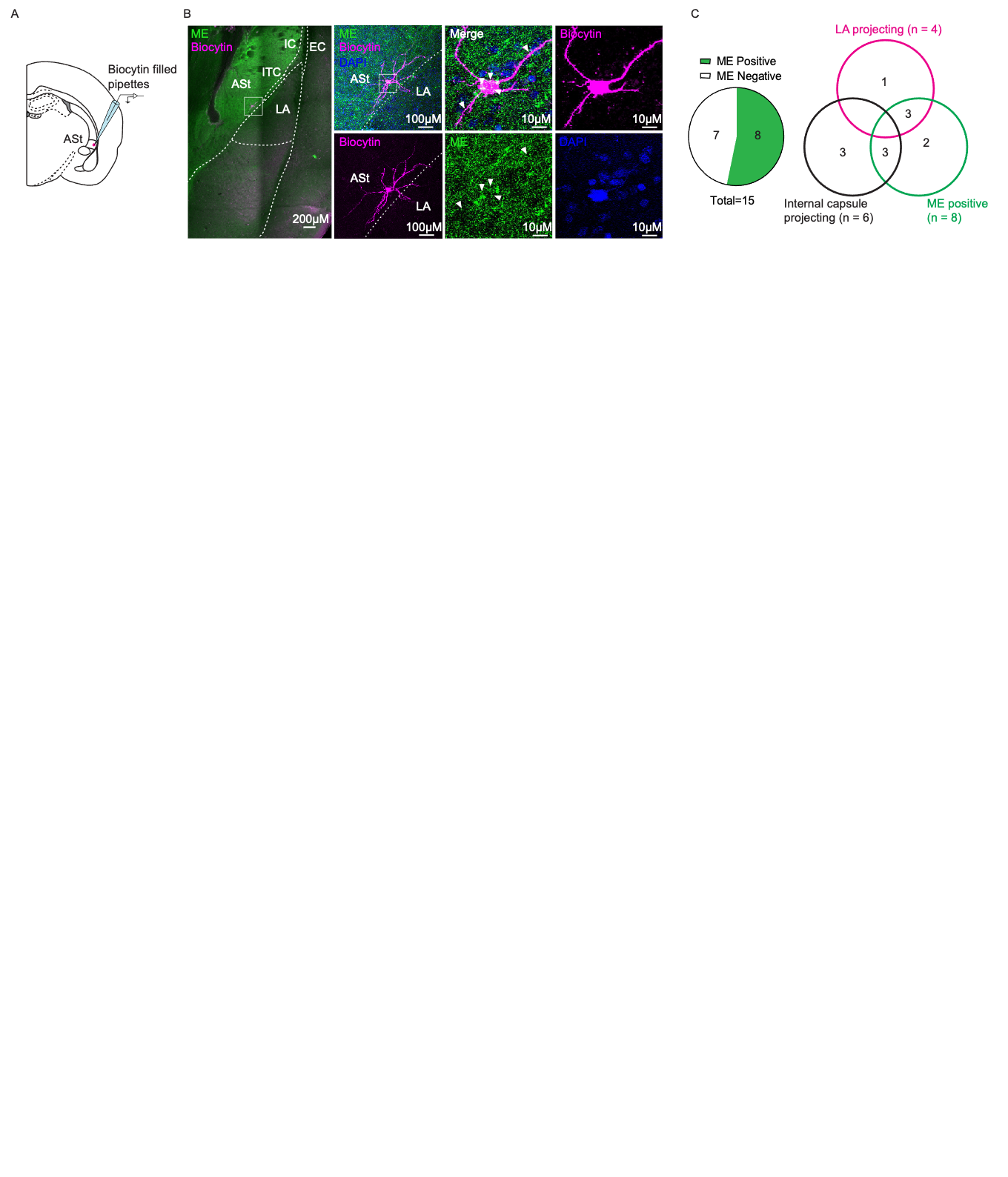

**(A)** Schematic diagram indicating recording locations for ASt MSNs. **(B)** *Post hoc* confocal images of met-enkephalin (green) and biocytin-labelled MSN (purple). Low power image shows met-enkephalin immunoreactivity around the biocytin-labelled cell. High powered magnification reveals dendritic spines and co-localisation of biocytin with met-enkephalin. **(C)** Left: pie chart indicating 8/15 biocytin-filled neurones were met-enkephalin positive. Right: Venn diagram indicating the proportion of met-enkephalin positive neurones and their dendritic projection targets (4/15 terminating in the LA, 6/15 terminating at the ASt-LA border).

**Supplementary Figure 4. PENK knockdown is specific to the injection site in the ASt**

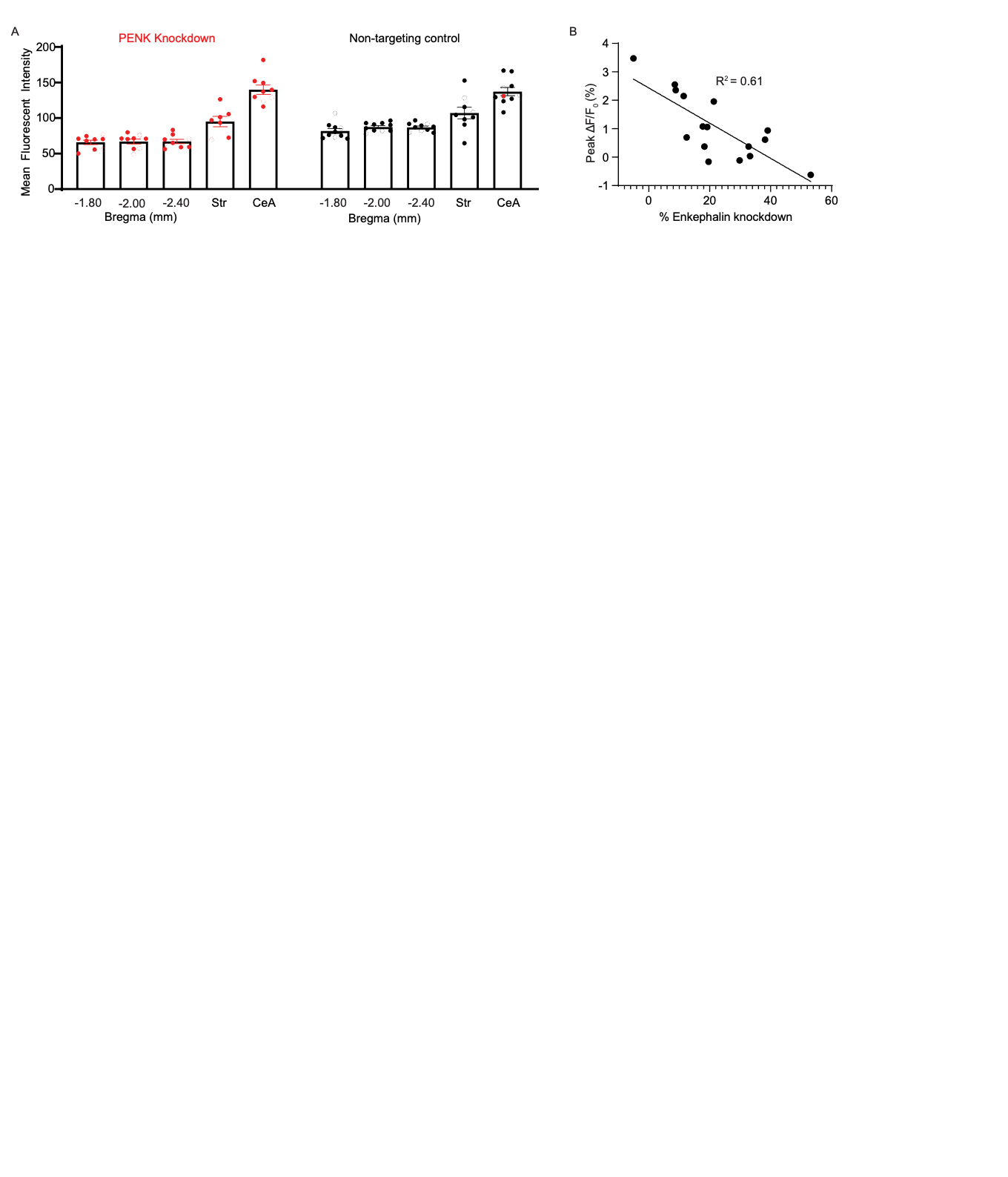

**(A**) Graph showing mean fluorescent intensity as measured in the ASt across the rostral-caudal range and in the striatum and CeA. **(B)** Correlation analysis between % of enkephalin knockdown and peak ΔF/F_0._ R^2^ = 0.61

**Supplementary Figure 5. Functional connection from the VTA to ASt are sparse.**

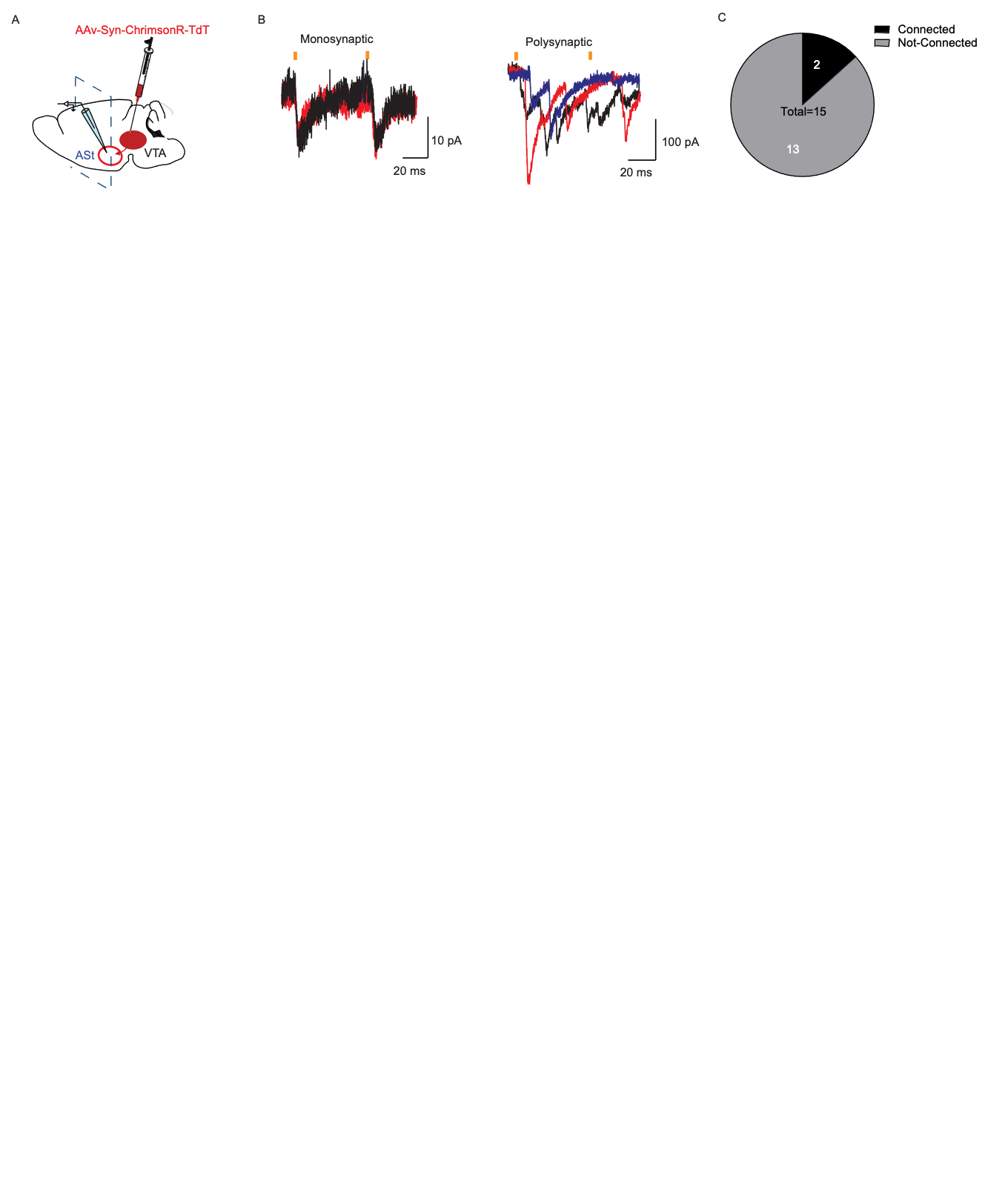

**(A)** Schematic diagram showing injection of ChrimsonR-TdT and recording site in the ASt. **(B)** Example traces showing oEPSC’s from putative ASt MSNs indicating a directly (left) and indirectly (right) connected cell. Black, red and blue traces indicate different stimulations episode from the same recording. Orange bars indicate optical stimulation points. **(C)** Pie chart showing number of connected verses non-connected cells. A total of 2 cells shown in (B) were connected our of 15 neurons recorded

**Supplementary Figure 6. Injections sites for ChrimsonR-TDT in the VTA**

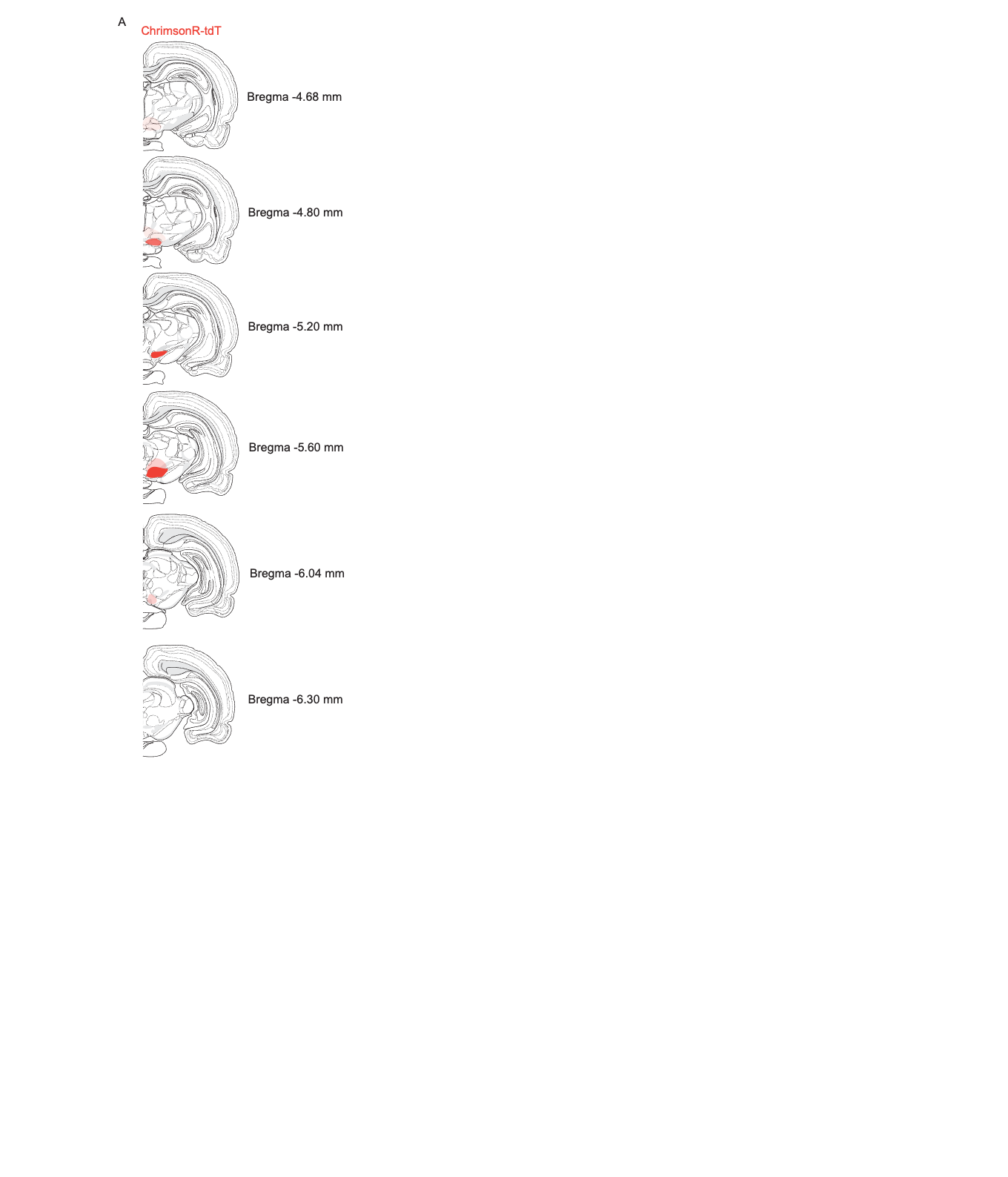

**(A**) Schematic diagram showing injection sites of ChrimsonR-TdT injection at various rostral-caudal slices of the VTA. Dark colours indicate regions of high cell body expression.
